# Immunization with recombinant accessory protein-deficient SARS-CoV-2 protects against lethal challenge and viral transmission

**DOI:** 10.1101/2022.03.13.484172

**Authors:** Chengjin Ye, Jun-Gyu Park, Kevin Chiem, Piyush Dravid, Anna Allué-Guardia, Andreu Garcia-Vilanova, Amit Kapoor, Mark R. Walter, James J. Kobie, Richard K. Plemper, Jordi B. Torrelles, Luis Martinez-Sobrido

## Abstract

Severe Acute Respiratory Syndrome Coronavirus 2 (SARS-CoV-2) has led to a worldwide Coronavirus Disease 2019 (COVID-19) pandemic. Despite high efficacy of the authorized vaccines, protection against the surging variants of concern (VoC) was less robust. Live-attenuated vaccines (LAV) have been shown to elicit robust and long-term protection by induction of host innate and adaptive immune responses. We sought to develop a COVID-19 LAV by generating 3 double open reading frame (ORF)-deficient recombinant (r)SARS-CoV-2 simultaneously lacking two accessory open reading frame (ORF) proteins (ORF3a/ORF6, ORF3a/ORF7a, and ORF3a/ORF7b). Here, we report that these double ORF-deficient rSARS-CoV-2 have slower replication kinetics and reduced fitness in cultured cells as compared to their parental wild-type (WT) counterpart. Importantly, these double ORF-deficient rSARS-CoV-2 showed attenuation in both K18 hACE2 transgenic mice and golden Syrian hamsters. A single intranasal dose vaccination induced high levels of neutralizing antibodies against different SARS-CoV-2 VoC, and also activated viral component-specific T-cell responses. Notably, the double ORF-deficient rSARS-CoV-2 were able to protect, as determined by inhibition of viral replication, shedding, and transmission, against challenge with SARS-CoV-2. Collectively, our results demonstrate the feasibility to implement these double ORF-deficient rSARS-CoV-2 as safe, stable, immunogenic and protective LAV for the prevention of SARS-CoV-2 infection and associated COVID-19 disease.

## INTRODUCTION

Coronaviruses (CoV) cause mild to lethal respiratory infections ^1,2^. Severe Acute Respiratory Syndrome CoV 2 (SARS-CoV-2) is the etiological agent of the worldwide CoV Disease 2019 (COVID-19) pandemic, which began at the end of 2019 in the city of Wuhan in China. As of March 2022, more than 450 million cases and over 6 million deaths have been confirmed worldwide (https://COVID-19.who.int/). SARS-CoV-2 belongs to the *Coronaviridae* family, order *Nidovirales*, and is an enveloped positive-sense, single-stranded RNA virus with a large and non-segmented genome of ∼30 kb in length. The SARS-CoV-2 genome encodes a major open reading frame (ORF) frameshifted polyprotein (ORF1a/ORF1ab) and four structural proteins: the spike (S), envelope (E), membrane (M) and nucleocapsid (N) proteins ^3^. In addition, at least six ORFs encoding accessory proteins are interspersed between the structural genes ^4^. These accessory proteins have been reported to regulate different aspects of the host response to viral infection, including cell apoptosis, interferon (IFN) signaling, and immune modulation, among others ^5-9^.

Subunit-, inactivated-, mRNA-, and vector-based vaccines have been developed for SARS-CoV-2, and the United States Food and Drug Administration (FDA) has authorized the clinical use of 2 types of mRNA vaccines and 1 adenoviral vector vaccine. These vaccines rely on the expression of the viral S protein to induce high levels of neutralizing antibodies against SARS-CoV-2 ^10,11^. However, when SARS-CoV-2 variants of concern (VoC) emerged ^12,13^, the efficacy of these vaccines significantly declined because of the narrow plasticity of the immune response against the S protein. Vaccines that induce comprehensive immunity against multiple vial components, in addition to the viral S glycoprotein, promise to provide enhanced potency to diverse VoC to fight the ongoing COVID-19 pandemic. Live-attenuated vaccines (LAV) have traditionally been highly successful at inducing effective immunity. Since LAV most closely resemble a natural infection, yet have diminished pathogenesis, a comprehensive immune response is induced, calling on B cell and T cell responses to not only target viral glycoproteins but also other viral components. For example, the Yellow Fever LAV is considered one of the safest and most efficacious vaccines developed to date ^14-16^. Initial attempts to generate a SARS-CoV-2 LAV using a mutant virus that lacked the furin cleavage site in the S protein, showed reduced viral pathogenesis in both K18 hACE2 transgenic mice and golden Syrian hamsters, and conferred protection against challenge with the parental virus ^17^. Another intranasal LAV strategy used a highly attenuated SARS-CoV-2 that incorporated a furin deletion plus a S protein segment with synonymous suboptimal codon pairs (codon-pair deoptimization), which protected hamsters from SARS-CoV-2-associated weight loss ^18^. Additionally, the codon-pair deoptimization strategy was extended to the entire viral genome, resulting in two recombinant (r)SARS-CoV-2 LAV candidates that induced strong protective immunity in hamsters, blocking all clinical symptoms ^19^. However, the attenuated SARS-CoV-2 generated in these studies as LAV still have a high risk to revert, and their efficacies, including against infection and transmission, have not been fully evaluated.

Previously, we developed rSARS-CoV-2 by deleting single ORF viral accessory proteins (ORF3a, ORF6, ORF7a, ORF7b, and ORF8) ^20^. However, these single ORF-deficient rSARS-CoV-2 were still lethal in the K18 hACE2 transgenic mouse model, with only the rSARS-CoV-2 ΔORF3a showing attenuation, with a 75% survival rate ^20^. In this study, we use our previously described reverse genetics system ^21^ to extend the attenuation strategy by starting with rSARS-CoV-2 ΔORF3a to generate recombinant viruses with a second ORF deletion (ORF3a/ORF6, ORF3a/ORF7a, ORF3a/ORF7b, and ORF3a/ORF8). While the double ORF-deficient rSARS-CoV-2 Δ3a/ Δ 8 did not result in a viable virus, the rSARS-CoV-2 harboring deletions of ORF3a/ORF6 (Δ3a/Δ6), ORF3a/ORF7a (Δ3a/Δ7a), or ORF3a/ORF7b (Δ3a/Δ7b) were successfully rescued and demonstrated slower kinetics and reduced fitness in cultured cells. Survival of K18 hACE2 mice infected with either of the 3 double ORF-deficient rSARS-CoV-2 was significantly increased. However, their survival rate varied from 0% for rSARS-CoV-2 Δ3a/Δ7a, 60% for rSARS-CoV-2 Δ3a/Δ6, to 100% for rSARS-CoV-2 Δ3a/Δ7b. Notably, both humoral and cellular immunity were activated in the surviving K18 hACE2 transgenic mice to protect them against subsequent lethal challenge with parental SARS-CoV-2. Similarly, hamsters vaccinated with the 3 double ORF-deficient rSARS-CoV-2 mounted robust levels of neutralizing antibodies without leading to any body weight changes. Importantly, upon challenge with parental SARS-CoV-2, all vaccinated hamsters showed decreased viral replication, shedding and transmission.

Together, our results demonstrate that the double ORF-deficient rSARS-CoV-2 developed in this study are promising candidates for their implementation as safe, stable, immunogenic, and protective LAV for the prophylactic treatment of SARS-CoV-2 infection and associated COVID-19 disease.

## RESULTS

### Generation and *in vitro* characterization of rSARS-CoV-2 with deletions of two accessory proteins

We generated 3 double ORF-deficient rSARS-CoV-2 that simultaneously lack two accessory proteins (Δ3a/Δ6, Δ3a/Δ7a, and Δ3a/Δ7b) using a previously described bacterial artificial chromosome (BAC)-based reverse genetic system ^21^ (**Extended Data Fig. 1A**). The 3 double ORF-deficient rSARS-CoV-2 were verified by RT-PCR amplification of the ORF3a, ORF6, ORF7a, ORF7b, and N viral genes (**Extended Data Fig. 1B**), and next generation sequencing of the viral genomes. In total, four non-reference alleles were found in these viruses with a frequency greater than 10% in corresponding populations: two non-reference alleles (S N74K and ORF8 S21G) were found in rSARS-CoV-2 Δ3a/Δ6 with a percentage of 58.52% and 57.69%, respectively (**Extended Data Fig. 1C, top**), one non-reference allele (ORF8 A51G) in rSARS-CoV-2 Δ3a/Δ7a with a percentage of 12.04% (**Extended Data Fig. 1C, middle**), and a non-reference allele (E L37H) in rSARS-CoV-2 Δ3a/Δ7b with a percentage of 14.94% (**Extended Data Fig. 1C, bottom**).

Having validated the double ORF-deficient rSARS-CoV-2, we evaluated their plaque morphology in Vero E6 cells at 24, 48, 72, and 96 hours post-infection (hpi) (**Fig. 1A**). All 3 double ORF-deficient rSARS-CoV-2 exhibited smaller plaque diameters at 72 and 96 hpi than rSARS-CoV-2 WT (**Fig. 1B**). Then, we evaluated the plaque phenotype stability of these double ORF-deficient rSARS-CoV-2 after 10 serial passages in Vero E6 cells. Passage 10 (P10) double ORF-deficient rSARS-CoV-2 showed a similar plaque phenotype with comparable plaque size diameters to those of P1 (**Fig. 1C**). Lastly, growth kinetics of these double ORF-deficient rSARS-CoV-2 were evaluated in Vero E6 and A549 expressing human ACE2 (A549 hACE2) cells infected with a multiplicity of infection (MOI) of 0.01 plaque-forming units (PFU) per cell. All double ORF-deficient rSARS-CoV-2 replicated to similar peak titers, which were significantly lower than those reached by rSARS-CoV-2 WT at 24, 48, 72 and 96 hpi in both cell lines (**Fig. 1D**).

**Figure 1.**
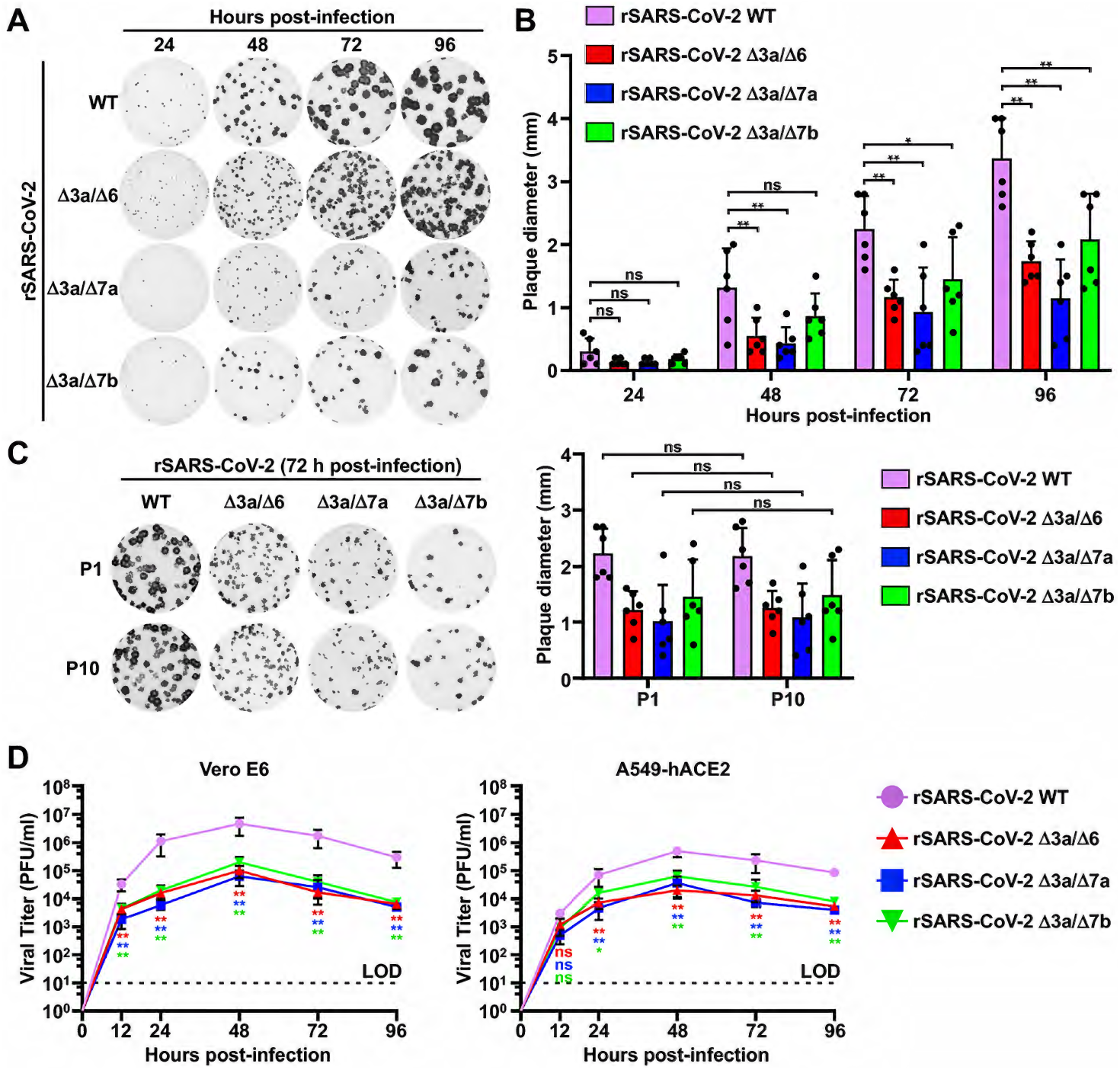
*In vitro* characterization of the double ORF-deficient rSARS-CoV-2. **(A)** Plaque phenotype. Plaque phenotypes of the WT and double ORF-deficient rSARS-CoV-2 in Vero E6 cells. Plaques were visualized by immunostaining with a monoclonal antibody (1C7C7) against the viral N protein. **(B)** Viral plaque size analysis. Six plaques were randomly selected and measured using a standard ruler (millimeters, mm). Data is presented as means ± standard deviation (SD). *, *P*<0.05; **, *P*<0.01; ns, not significant. **(C)** *In vitro* stability. The P1 and P10 of the WT and double ORF-deficient rSARS-CoV-2 were analyzed by plaque assay in Vero E6 cells. The plaques were visualized by immunostaining with the 1C7C7 monoclonal antibody against the viral N protein, and six plaques were random selected and measured with a standard ruler (mm). Data is presented as means ± SD. ns, not significant. **(D)** Growth kinetics. Viral growth kinetics of the WT and double ORF-deficient rSARS-CoV-2 in Vero E6 (left) and A549-hACE2 (right) cells. Dotted lines indicate the limit of detection (LOD). Data is presented as means ± SD. Statistical analysis was performed between the double ORF-deficient rSARS-CoV-2 and rSARS-CoV-2 WT. *, *P*<0.05; **, *P*<0.01; ns, not significant.

### Characterization of double ORF-deficient rSARS-CoV-2 in K18 hACE2 transgenic mice

We next evaluated replication and pathogenesis of the double ORF-deficient rSARS-CoV-2 in transgenic mice that express human angiotensin converting enzyme 2 (K18 hACE2), which are a common model to study SARS-CoV-2 pathogenesis ^22,23^. To that end, five-week-old female K18 hACE2 transgenic mice were infected with the double ORF-deficient rSARS-CoV-2 using a dose of 2×10^5^ PFU/mouse (**Fig. 2A**). Mock-infected K18 hACE2 mice and mice infected with the same dose of rSARS-CoV2 WT were used as internal controls. The lungs of infected K18 hACE2 transgenic mice were excised, and gross pathological lesions were analyzed at 2 and 4 days post-infection (dpi) (**Fig. 2B**). At 2 dpi, the lungs of K18 hACE2 transgenic mice infected with rSARS-CoV-2 WT contained pathological lesions in ∼35% of the total lung area, whereas the lungs of K18 hACE2 transgenic mice infected with the double ORF-deficient rSARS-CoV-2 had lesions in ≤ 25% of the total lung area with 19% for rSARS-CoV-2 Δ3a/Δ6, 23% for rSARS-CoV-2 Δ3a/Δ7a and 12% for rSARS-CoV-2 Δ3a/Δ7b. By 4 dpi, rSARS-CoV-2 WT had induced lesions in ∼60% of the total lung area, while double ORF-deficient rSARS-CoV-2 had reduced lesions with ∼40% for rSARS-CoV-2 Δ3a/Δ6, ∼55% for rSARS-CoV-2 Δ3a/Δ7a, and ∼40% for rSARS-CoV-2 Δ3a/Δ7b (**Fig. 2C**). We also evaluated viral replication of the double ORF-deficient rSARS-CoV-2 in the lungs and nasal turbinates of the infected K18 hACE2 transgenic mice. Compared with rSARS-CoV-2 WT, all double ORF-deficient rSARS-CoV-2 replicated to significantly lower titers in the lungs and nasal turbinates of infected mice at both 2 and 4 dpi (**Fig. 2D**). We further analyzed Th1/Th2/Th17 immune responses and chemokines present in the lungs of K18 hACE2 mice infected with the double ORF-deficient rSARS-CoV-2 (**Extended Data Fig. 2A**). At 2 dpi, there were no major differences among all virus strains studied, except for the rSARS-CoV-2 Δ3a/Δ7a-infected samples, which showed significantly lower levels of IFN responses (IFN-α and IFN-γ). However, by 4 dpi, all the samples from the double ORF-deficient rSARS-CoV-2 infected mice showed a decrease in inducing IFN-α when compared to that of rSARS-CoV-2 WT, and the decrease was not observed for IFN-γ. All the double ORF-deficient rSARS-CoV-2 infection also induced significantly lower levels of chemokines by 4 dpi.

**Figure 2.**
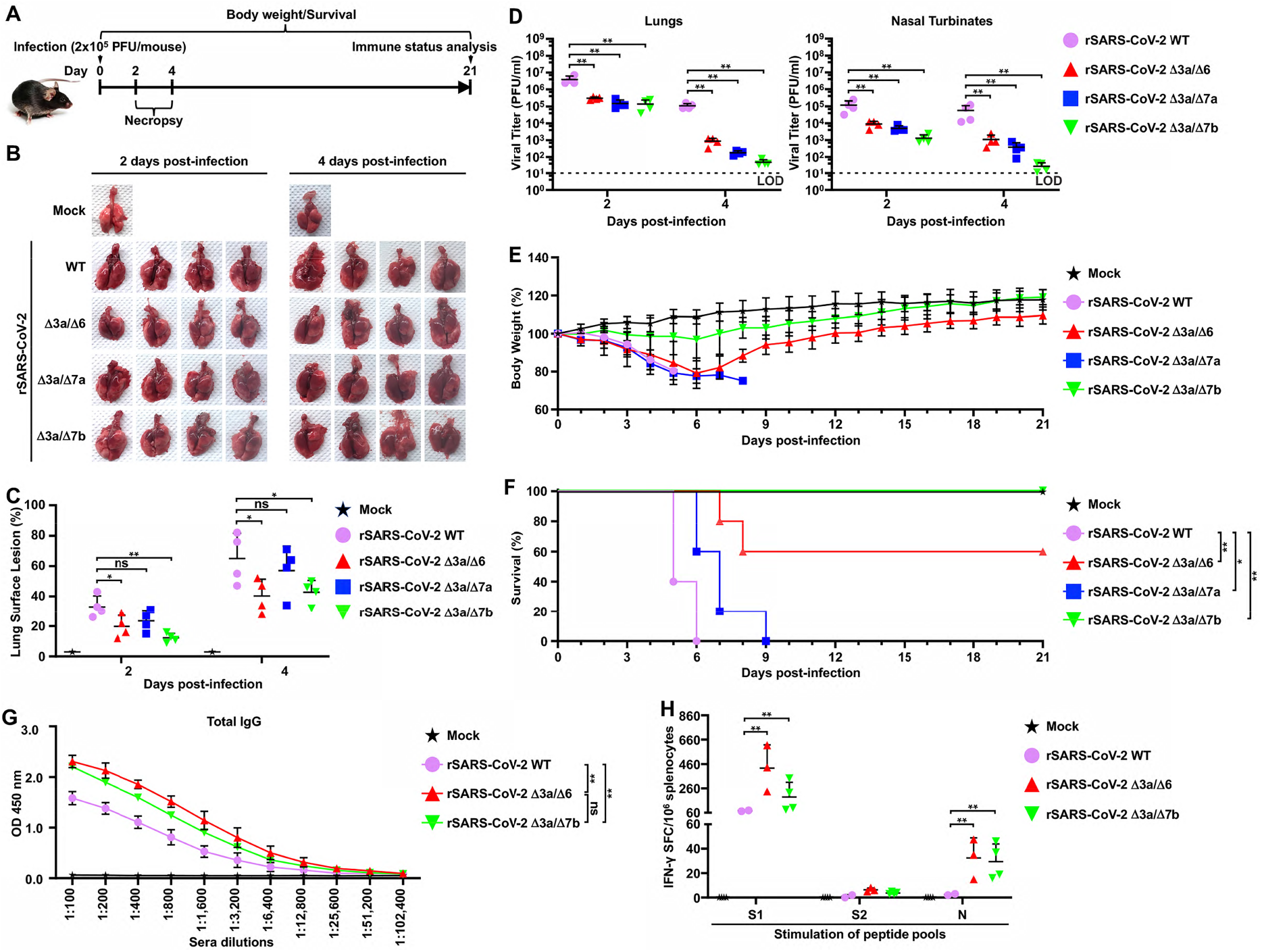
Characterization of double ORF-deficient rSARS-CoV-2 in K18 hACE2 transgenic mice. **(A)** Schematic representation of the experimental timeline used to infect K18 hACE2 transgenic mice with the WT and double ORF-deficient rSARS-CoV-2. **(B)** Pathological lesions in the lung surface of K18 hACE2 transgenic mice mock-infected or infected (2×10^5^ PFU/mouse, n=4) with the indicated rSARS-CoV-2 at 2 and 4 dpi. **(C)** Pathological lesion scoring on the lung images in panel B using ImageJ. **(D)** Viral titers in the clarified homogenate of lungs (left) and nasal turbinates (right) of K18 hACE2 transgenic mice infected in panel B at 2 and 4 dpi. **(E)** Body weight changes in K18 hACE2 transgenic mice mock-infected or infected (2×10^5^ PFU/mouse, n=5) with the indicated WT or double ORF-deficient rSARS-CoV-2. **(F)** Survival curves of K18 hACE2 transgenic mice infected in E were calculated and plotted using daily observations for 21 days. *, *P*<0.05; **, *P*<0.01. **(G)** Total levels of IgG against viral full-length S protein in the sera from the mice that survived in panel F were tested by ELISA at 21 dpi. The sera collected from the two surviving K18 hACE2 mice infected with rSARS-CoV-2 WT (10^3^ PFU/mouse, n=5) for 21 days were included as a positive control. **, *P*<0.01; ns, not significant. **(H)** Splenocytes were isolated from the mice that survived in panel F at 21 dpi, and IFN-γ-specific spot-forming cells (SFC) were counted after stimulation with peptide pools of S1, S2, and N using flow cytometry. The splenocytes isolated from the two surviving K18 hACE2 mice infected with rSARS-CoV-2 WT (10^3^ PFU/mouse, n=5) for 21 days were included as a positive control. **, *P*<0.01.

Clinical signs demonstrated that K18 hACE2 transgenic mice infected with rSARS-CoV-2 WT started losing bodyweight at 2 dpi and succumbed to infection by 6 dpi, and a comparable pattern of bodyweight loss was observed in animals infected with rSARS-CoV-2 Δ3a/Δ6 and rSARS-CoV-2 Δ3a/Δ7a (**Fig. 2E**). All K18 hACE2 transgenic mice infected with rSARS-CoV-2 Δ3a/Δ7a and 2 out of the 5 mice infected with rSARS-CoV-2 Δ3a/Δ6 succumbed to infection by 9 dpi infection (**Fig. 2F**). The other 3 K18 hACE2 transgenic mice infected with rSARS-CoV-2 Δ3a/Δ6 gradually recovered and ultimately survived from viral infection (**Figs. 2E and 2F**). Notably, all K18 hACE2 transgenic mice infected with rSARS-CoV-2 Δ3a/Δ7b maintained their initial body weight, with only 2 out of the 5 mice losing a maximum of ∼15% of their initial bodyweight before 6 dpi (**Fig. 2E**). Mortality analysis showed a survival rate of 60% for K18 hACE2 transgenic mice infected with rSARS-CoV-2 Δ3a/Δ6 and 100% survival rate for K18 hACE2 transgenic mice infected with rSARS-CoV-2 Δ3a/Δ7b. All mice infected with rSARS-CoV-2 Δ3a/Δ7a succumbed to viral infection, similar to mice infected with rSARS-CoV-2 WT, although the survival time in mice infected with rSARS-CoV-2 Δ3a/Δ7a was significant increased compared to mice infected with rSARS-CoV-2 WT (**Fig. 2F**). At 21 dpi, high levels of immunoglobulin G (IgG) against full-length viral S protein were detected in the sera of surviving mice **(Fig. 2G**), which are higher than that detected in the sera from K18 hACE2 transgenic mice infected with a sublethal dose of 10^3^ PFU/mouse rSARS-CoV-2 WT (**Fig. 2G**). Importantly, viral Spike 1 (S1), Spike 2 (S2) and N protein-specific IFN-γ secreting cells were detected in the splenocytes of surviving mice (**Fig. 2H**), and viral S1, E and M protein-specific cytokine-positive CD4^+^ and CD8^+^ T cells were also identified in both rSARS-CoV-2 Δ3a/Δ6 and rSARS-CoV-2 Δ3a/Δ7a infected K18 hACE2 transgenic mice splenocytes (**Extended Data Figs. 2B** and **2C**).

### Protection of rSARS-CoV-2 Δ3a/Δ7b-vaccinated K18 hACE2 transgenic mice against lethal challenge with SARS-CoV-2

Since infection with rSARS-CoV-2 Δ3a/Δ7b was not lethal in K18 hACE2 transgenic mice (**Figs. 2E** and **2F**), but induced robust humoral (**Fig. 2G**) and cellular (**Fig. 2H**) immunity, we hypothesized that the K18 hACE2 transgenic mice vaccinated with rSARS-CoV-2 Δ3a/Δ7b would survive a lethal challenge with wild-type virus, confirming the feasibility of rSARS-CoV-2 Δ3a/Δ7b as a LAV. To test this hypothesis, five-week-old female K18 hACE2 transgenic mice were either mock-vaccinated or rSARS-CoV-2 Δ3a/Δ7b-vaccinated with 2×10^5^ PFU/mouse; and intranasally challenged with 10^5^ PFU/mouse of rSARS-CoV-2 mCherryNluc at 21 days post-vaccination (**Fig. 3A**). Viral replication was evaluated using a non-invasive *in vivo* imaging system (IVIS) in the whole organism (Nluc), as previously described ^24^. Strong Nluc signal in the lungs of mock-vaccinated mice was detected at 2 and 4 days post-challenge with rSARS-CoV-2 mCherryNluc, whereas no Nluc signal was detected in the lungs of K18 hACE2 transgenic mice vaccinated with rSARS-CoV-2 Δ3a/Δ7b (**Figs. 3B** and **3C**). In excised lungs, expression of mCherry was readily detected in mock-vaccinated mice, while no mCherry was expressed in the lungs of mice vaccinated with rSARS-CoV-2 Δ3a/Δ7b (**Extended Data Figs. 3A** and **3B**). Moreover, pathological lesions on the lung surface were much more severe in mock-vaccinated mice compared to those in rSARS-CoV-2 Δ3a/Δ7b-vaccinated mice (**Extended Data Figs. 3C** and **3D**). Supernatants of the lung and nasal turbinate homogenates from rSARS-CoV-2 Δ3a/Δ7b-vaccinated mice did not contain infectious virus at both 2 and 4 days post-challenge (**Extended Data Fig. 3E**). Significantly decreased Nluc activity was further seen in the supernatants of the lung and nasal turbinate homogenates from rSARS-CoV-2 Δ3a/Δ7b-vaccinated K18 hACE2 transgenic mice at both 2 and 4 days post-challenge compared to mock-vaccinate mice (**Extended Data Fig. 3F**), which correlated with the *in vivo* imaging data. In the lungs of the mock-vaccinated mice, a significant production of IFN-α and IFN-γ were induced after challenge of rSARS-CoV-2 mCherryNluc (**Extended Data Fig. 3G**). In contrast, the IFN responses were not induced in the lungs of rSARS-CoV-2 Δ3a/Δ7b-vaccinated mice at either 2 or 4 days post-challenge, whereas an elevated production of TNF-α, which is highly related to a protective Th17 response, was induced at 4 days post-challenge (**Extended Data Fig. 3G**).

**Figure 3.**
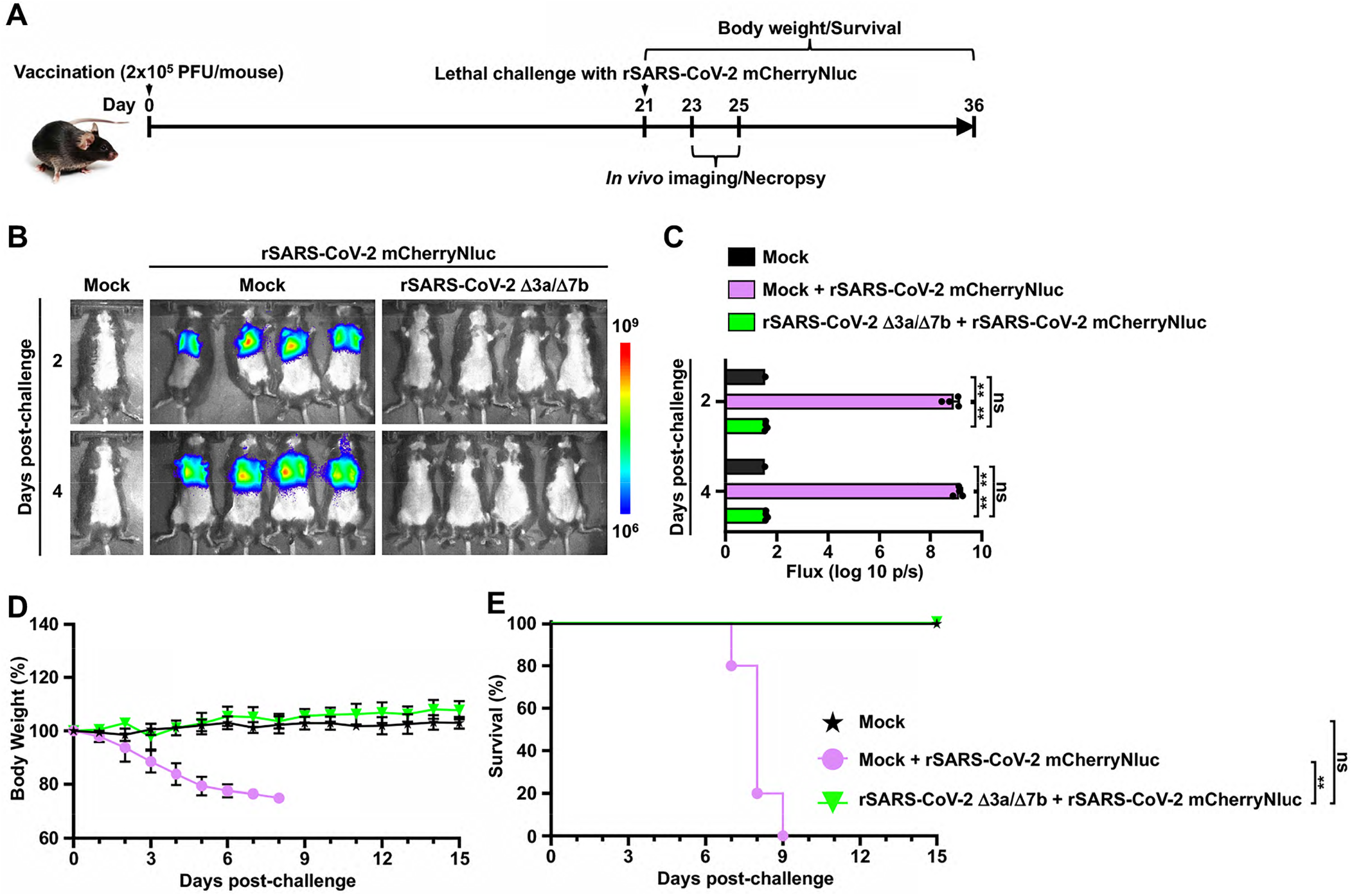
Protection efficacy of rSARS-CoV-2 Δ3a/Δ7b-vaccinated K18 hACE2 transgenic mice against lethal challenge with SARS-CoV-2. **(A)** Schematic representation of the experimental timeline used for the protection studies with rSARS-CoV-2 Δ3a/Δ7b in K18 hACE2 transgenic mice challenge with rSARS-CoV-2 mCherryNluc. **(B)** *In vivo* imaging of K18 hACE2 transgenic mice mock-vaccinated or vaccinated with rSARS-CoV-2 Δ3a/Δ7b at 2 and 4 days post-challenge with rSARS-CoV-2 mCherryNluc (n=4). Mock-vaccinated and mock-challenge K18 hACE2 transgenic mice were used as controls. **(C)** Quantitative analysis of Nluc expression in K18 hACE2 transgenic mice from B. **, *P*<0.01; ns, not significant. **(D)** Body weight changes of mock-vaccinated or rSARS-CoV-2 Δ3a/Δ7b-vaccinated K18 hACE2 transgenic mice were monitored for 15 days after challenge with rSARS-CoV-2 mCherryNluc. Mock-vaccinated and mock-challenge K18 hACE2 transgenic mice were used as controls. **(E)** Survival curves of mock-vaccinated or rSARS-CoV-2 Δ3a/Δ7b-vaccinated K18 hACE2 transgenic mice after challenge with rSARS-CoV-2 mCherryNluc. Mock-vaccinated and mock-challenge K18 hACE2 transgenic mice were used as controls. **, *P*<0.01; ns, not significant.

As expected, mock-vaccinated K18 hACE2 transgenic mice lose body weight starting at 2 days post-challenge (**Fig. 3D**), and all of them succumbed to infection by day 9 post-challenge with rSARS-CoV-2 mCherryNluc (**Fig. 3E**). In contrast, all mice vaccinated with rSARS-CoV-2 Δ3a/Δ7b showed no clinical signs of disease, including changes in body weight, and all mice (100%) survived the lethal challenge with rSARS-CoV-2 mCherryNluc (**Figs. 3D** and **3E**).

### Double ORF-deficient rSARS-CoV-2 vaccination prevents viral replication and shedding in hamsters

Next, we sought to investigate the safety and protective efficacy of the double ORF-deficient rSARS-CoV-2 in hamsters, an animal model of SARS-CoV-2 pneumonia that better recapitulates human COVID-19 than K18 hACE2 mice ^21^. Five-week-old female golden Syrian hamsters were mock-vaccinated or vaccinated (4×10^5^ PFU) with the double ORF-deficient rSARS-CoV-2 and monitored daily for body weight. After 21 days, hamsters were challenged with 2×10^5^ PFU of rSARS-CoV-2 mCherryNluc. To assess viral shedding and transmission, each challenge hamster (donor) was housed in the same cage with a susceptible contact hamster at 1 day post-challenge (**Fig. 4A**). Infection with rSARS-CoV-2 WT led to an ∼15% body weight loss by 6 dpi, and no changes in body weight were observed in hamsters infected with these double ORF-deficient rSARS-CoV-2, whose body weight were comparable to mock-infected animals at all time points (**Fig. 4B**). After viral challenge, the Nluc signal in all donor and contact hamsters were determined at 2 and 4 days post-challenge. In the donor hamsters, Nluc signal was detected in the nasal turbinates and lungs of mock-vaccinated hamsters at both time points; to a lesser extent, Nluc signal was also present in nasal turbinates, but not lungs, of the double ORF-deficient rSARS-CoV-2 vaccinated hamsters at 2 days post-challenge (**Figs. 4C** and **4D**). No Nluc was detected in all of these hamsters at 4 days post-challenge (**Figs. 4C** and **4D**). In contact hamsters, the Nluc signal was absent in any contact hamsters at 2 and 4 days post-challenge, but readily detectable in all hamsters in contact with mock-vaccinated hamsters at 4 days post-challenge (**Figs. 4C** and **4D**).

**Figure 4.**
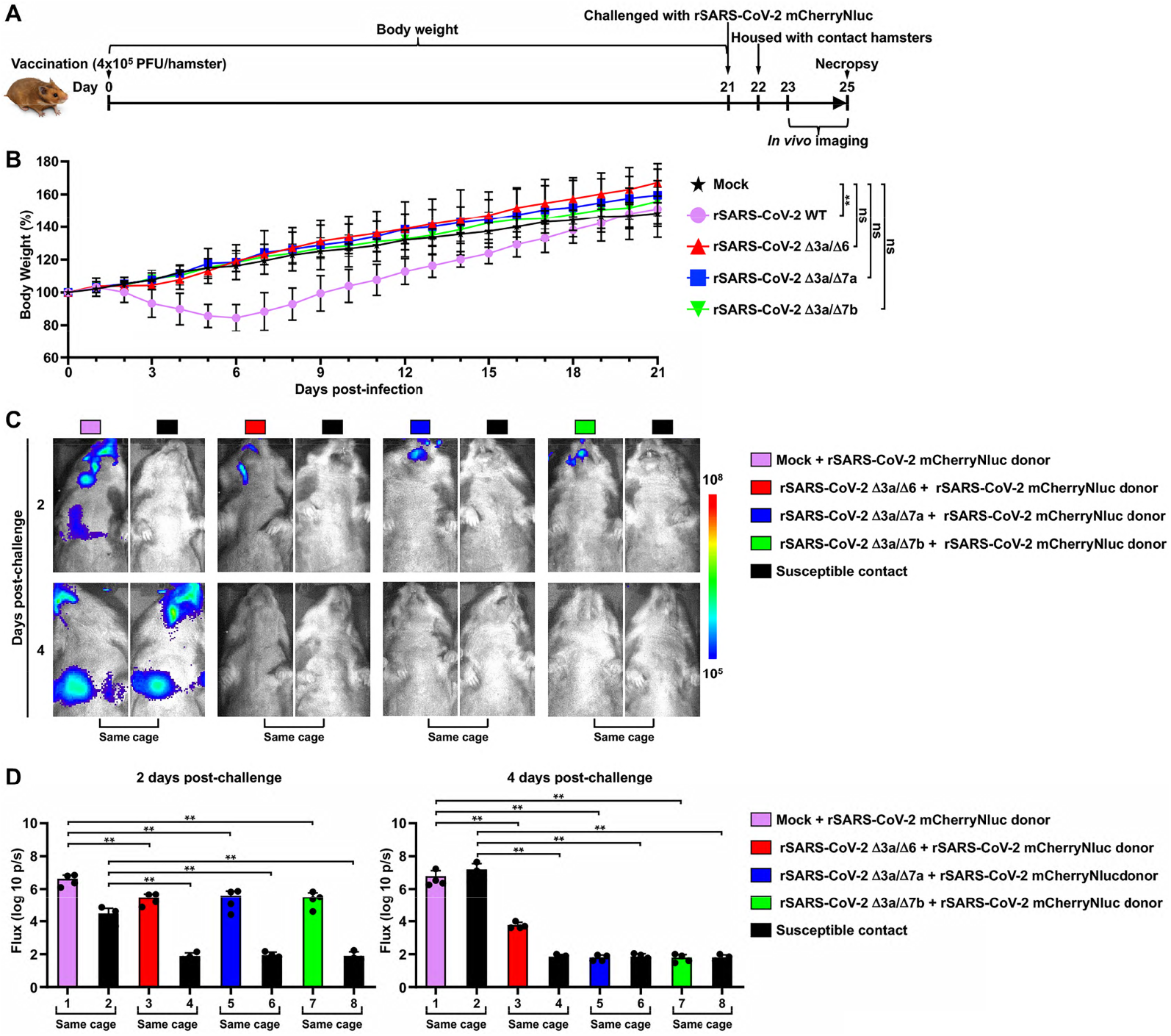
Double ORF-deficient rSARS-CoV-2 vaccination prevents replication and shedding of SARS-CoV-2 in hamsters. **(A)** Schematic representation of the experimental timeline used for the protection studies with the double ORF-deficient rSARS-CoV-2 in hamsters. **(B)** Body weight changes of mock-infected and WT or double ORF-deficient rSARS-CoV-2 infected hamsters were monitored for 21 days. **, *P*<0.01; ns, not significant. **(C)** *In vivo* imaging of the rSARS-CoV-2 mCherryNluc replication in hamsters at 2 and 4 days post-challenge. **(D)** Quantitative analysis of Nluc expression in hamsters from panel C at 2 (left) and 4 (right) days post-challenge with rSARS-CoV-2 mCherryNluc.

After collecting lungs at 4 days post-challenge, we noted strong mCherry expression in the lungs of mock-vaccinated and rSARS-CoV-2 mCherryNluc-infected donor hamsters and their contacts, whereas mCherry fluorescence was significantly decreased in all vaccinated donor hamsters and their respective contacts (**Extended Data Figs. 4A** and **4B**). We next determined virus load in clarified supernatants of lung and nasal turbinate homogenates from both donor and contact hamsters. No detectable infectious virus was present in either tissue in any of the donor hamsters vaccinated with the double ORF-deficient rSARS-CoV-2 (**Extended Data Fig. 4C**). All contact hamsters were free of virus, except for one contact hamster (∼10^2^ PFU/ml) that was co-housed with a hamster vaccinated with rSARS-CoV-2 Δ3a/Δ6 (**Extended Data Fig. 4C**). We obtained equivalent results when following Nluc activity in the clarified supernatant of lung and nasal turbinate homogenates (**Extended Data Fig. 4D**).

### Double ORF-deficient rSARS-CoV-2 vaccination prevents transmission in hamsters

Since double ORF-deficient rSARS-CoV-2 vaccination significantly reduced viral replication and shedding of SARS-CoV-2 after challenge, we sought to explore whether the double ORF-deficient rSARS-CoV-2 can provide protection against viral transmission between vaccinated contact hamsters and infected donors. Five-week-old female golden Syrian hamsters were vaccinated with the double ORF-deficient rSARS-CoV-2 (4×10^5^ PFU), and sera were collected at 18 days post-vaccination. The vaccinated contact hamsters were housed with rSARS-CoV-2 mCherryNluc-infected donor hamsters at 21 days post-vaccination and all hamsters were analyzed by *in vivo* imaging and necropsy (**Fig. 5A**). Sera collected from the double ORF-deficient rSARS-CoV-2-vaccinated hamsters showed a high neutralizing potential against SARS-CoV-2 WA1 strain and different VoC (Alpha, α; Beta, β; Delta, δ; and Omicron, o) (**Figs. 5B** and **5C**). After co-housing rSARS-CoV-2 mCherryNluc-infected (2×10^5^ PFU) donor hamsters with the double ORF-deficient rSARS-CoV-2-vaccinated contact hamsters, Nluc signal was readily detected in all donor hamsters at 2 and 4 dpi, whereas no detectable Nluc signal was observed in any of the contact animals at 2 dpi. At 4 dpi, high levels of Nluc signal were present in all mock-vaccinated contact hamsters, but signal was extremely low in all double ORF-deficient rSARS-CoV-2-vaccinated contact hamsters (**Figs. 5D** and **5E**). In the lungs excised at 4 dpi, mCherry expression was readily detected in all donor hamsters and mock-vaccinated contact hamsters but not in any of double ORF-deficient rSARS-CoV-2-vaccinated contact hamsters (**Extended Data Figs. 5A** and **5B**). When titrating infectious particles in the clarified lung and nasal turbinate homogenates, no infectivity was present in tissues derived from any of the contacts co-housed with the hamsters vaccinated with the double ORF-deficient rSARS-CoV-2 (**Extended Data Fig. 5C**). Consistent with the viral titer results, Nluc activity in the clarified lung and nasal turbinate homogenates was significantly decreased in all double ORF-deficient rSARS-CoV-2-vaccinated contact hamsters compared to that present seen in mock-vaccinated contacts (**Extended Data Fig. 5D**).

**Figure 5.**
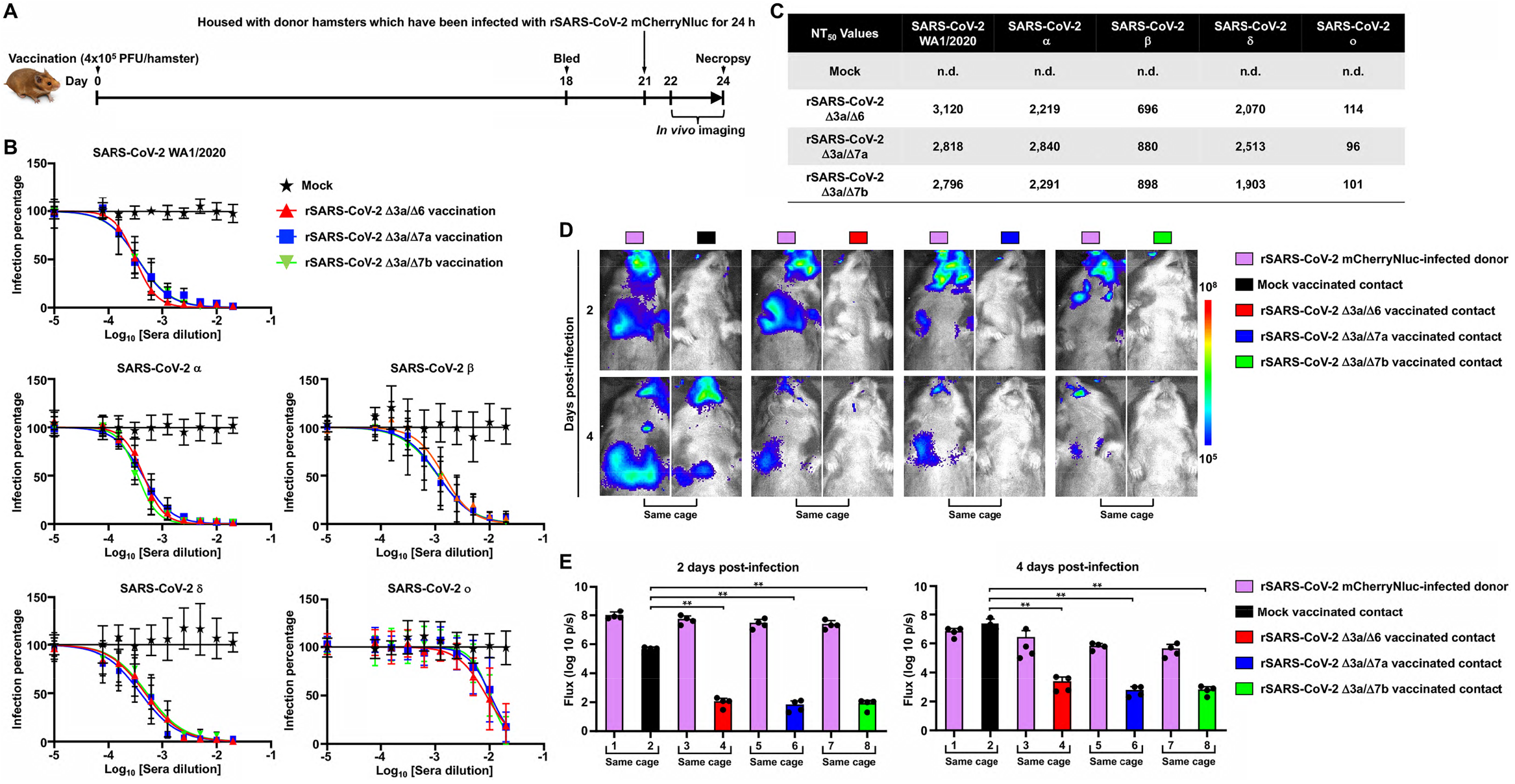
Double ORF-deficient rSARS-CoV-2 vaccination prevents transmission in hamsters. **(A)** Schematic representation for the experimental timeline used to test the prevention of transmission by double ORF-deficient rSARS-CoV-2 in hamsters. **(B)** Sera collected at 18 days post-vaccination were evaluated for the neutralizing capacity against SARS-CoV-2 WA1/2020, Alpha (α), Beta (β), Delta (δ), and Omicron (o) VoC by PRMNT assay. **(C)** Summary of NT_50_ values of sera against the different SARS-CoV-2 VoC. **(D)** *In vivo* imaging of the rSARS-CoV-2 mCherryNluc replication in hamsters at 2 and 4 dpi. **(E)** Quantitative analysis of Nluc expression in hamsters from panel D at 2 (left) and 4 (right) dpi.

## DISCUSSION

LAVs, which can induce broad host immune responses and activate both innate and adaptive immunity, are attractive for prophylaxis against emerging and re-emerging pathogens that undergoing rapid antigenic drift, such as SARS-CoV-2 ^25^. In this study, we used our established BAC-based reverse genetic system ^21^ to generate 3 potential SARS-CoV-2 LAV candidates by deleting two accessory protein ORFs simultaneously from the viral genome (ORF3a/ORF6, ORF3a/ORF7a, and ORF3a/ORF7b). After being passaged 10 times in Vero E6 cells, the 3 double ORF-deficient rSARS-CoV-2 showed similar plaque phenotypes as those found at P1, suggesting the genomes of the double ORF-deficient SARS-CoV-2 are stable *in vitro*. Results also showed that double ORF-deficient rSARS-CoV-2 grow efficiently in Vero E6 cells, a cell line approved by the United States FDA for vaccine manufacture and production.

K18 hACE2 transgenic mice were first developed for *in vivo* pathogenicity studies of SARS-CoV ^26^. Although K18 hACE2 transgenic mice display morbidity and mortality, including efficient replication in the upper and lower respiratory tract, and brain, upon SARS-CoV-2 infection ^27^, it is still unknown how much this K18 hACE2 transgenic mouse model recapitulate a true infection in humans. Since the natural isolate of one of the first SARS-CoV-2 WA1 strain exhibits a mouse lethal dose 50 (MLD_50_) of ∼500 PFU ^28^, we favored the use of the K18 hACE2 transgenic mouse model for testing the safety of our double ORF-deficient rSARS-CoV-2 LAV candidates. Using K18 hACE2 transgenic mice, we show our most attenuated double ORF-deficient virus, rSARS-CoV-2 Δ3a/Δ7b, has an MLD_50_ greater than 2×10^5^ PFU, and rSARS-CoV-2 Δ3a/Δ6 showed a MLD_50_ of ∼2×10^5^ PFU, which is about 400-fold higher than its WT counterpart, suggesting adequate attenuation of these two rSARS-CoV-2 Δ3a/Δ6 and rSARS-CoV-2 Δ3a/Δ7b LAV candidates. Although all mice infected with rSARS-CoV-2 Δ3a/Δ7b (2×10^5^ PFU) succumbed to infection, they still showed an increased survival time than K18 hACE2 transgenic mice infected with rSARS-CoV-2 WT. In addition, we noticed that pathological lesions were observed on the lungs of the 3 double ORF-deficient rSARS-CoV-2-infected K18 hACE2 mice at both 2 and 4 dpi, although reduced as compared to the pathological lesions observed in mice infected with rSARS-CoV-2 WT. However, viral replication of any LAV candidate was significantly reduced compared to that of rSARS-CoV-2 WT-infected K28 hACE2 transgenic mice.

Priming the immune system with one immunogen and then boosting with a different immunogen is known to enhance immunity of the host ^29,30^. Typically, the booster LAV is provided to a higher dose than the prime LAV so that the boost LAV is not neutralized by the immune response elicited by the prime vaccination ^29^. In this study, we have generated 3 SARS-CoV-2 LAV candidates, which have shown different virulence in K18 hACE2 transgenic mice, providing flexibility for designing different immunization protocols. Notably, in all cases, the 3 SARS-CoV-2 LAV candidates were more attenuated than a natural SARS-CoV-2 isolate, with rSARS-CoV-2 Δ3a/Δ7b being the most attenuated virus and rSARS-CoV-2 Δ3a/Δ7a the most virulent; and rSARS-CoV-2 Δ3a/Δ6 having an intermediate phenotype. It is worth noting that the K18 hACE2 transgenic mouse model is highly sensitive to SARS-CoV-2 infection and may not represent reflections observed in other animal models, or humans ^31-34^.

In the case of rSARS-CoV-2 Δ3a/Δ7b-infected K18 hACE2 transgenic mice, 2 of the 5-infected mice has some minimal body weight loss before 6 dpi. Of note, the fully attenuated rSARS-CoV-2 Δ3a/Δ7b was able to provide, upon a single intranasal administration, a protective effect in K18 hACE2 transgenic mice challenged with a lethal dose of rSARS-CoV-2 mCherryNluc, and 100% of the rSARS-CoV-2 Δ3a/Δ7b-vaccinated K18 hACE2 transgenic mice survived the challenge with SARS-CoV-2, suggesting that even the most attenuated LAV candidate vaccine could confer complete protection against a lethal challenge of SARS-CoV-2. Further, rSARS-CoV-2 Δ3a/Δ7b-vaccinated mice produced lower levels of IFN responses accompanied with significant decreases in chemokine production and a lower IL-6/IL-10 ratio. These results are consistent with an attenuated cytokine storm, which may explain the reduction in tissue damage observed and the better control of the infection by mice. The later Th1 response observed at 4 dpi indicate that rSARS-CoV-2 Δ3a/Δ7b-vaccinated mice might attenuate the rapid evolution of the detrimental cytokine response observed in the mock-vaccinated mice after challenge of rSARS-CoV-2 mCherryNluc.

Our hamster studies demonstrate that all 3 double ORF-deficient rSARS-CoV-2 are highly attenuated and infection with 4×10^5^ PFU does not result in body weight loss, contrary to hamsters infected with rSARS-CoV-2 WT, where animal lost up to 15% of their initial body weight by day 6, as previously described ^21^. Furthermore, virus replication in the double ORF-deficient rSARS-CoV-2-vaccinated hamsters after challenge was only transiently detected in the upper respiratory tract (nasal turbinate) at 2 but not at 4 dpi, which is very important for avoiding respiratory complications and pneumonia progression ^35^. Importantly, viral shedding was impeded in the double ORF-deficient rSARS-CoV-2-vaccinated hamsters as well, as suggested by a lack of viral replication at both 2 and 4 days post-challenge in susceptible contact hamsters. Moreover, vaccination with the double ORF-deficient rSARS-CoV-2 completely abrogated viral transmission from infected donor hamsters, as virus replication was completely undetectable in the vaccinated contact hamsters. The protection could be attributed to the high levels of neutralizing antibody found in sera, including neutralizing antibodies against different VoC, as well as an activated T-cell response. Interestingly, reduced virus replication was seen in infected donor hamsters, which were housed with double ORF-deficient rSARS-CoV-2-vaccinated contact hamsters. We speculate that the close proximity of the vaccinated contact hamster may allow the spread of viral-specific IgA through saliva, feces, and aerosols, which may neutralize virus replication in infected donors.

Together, our data show that the 3 double ORF-deficient rSARS-CoV-2, which express all viral structural proteins yet lack different accessory proteins, are attenuated to different degree. All double ORF-deficient rSARS-CoV-2 induce robust innate and adaptive immunogen responses upon a single intranasal administration that protects against subsequent challenge with SARS-CoV-2. Combining stable attenuation with high immunogenicity makes them an attractive option for their use as safe, stable, and protective LAV candidates for prophylaxis of SARS-CoV-2 infection and associated COVI-19 disease. This conclusion is based on comprehensive assessment in two different animal models of SARS-CoV-2 infection and transmission, the K18 hACE2 transgenic mouse ^22,23^ and the golden Syrian hamster models ^21^, respectively.

## MATERIALS AND METHODS

### Biosafety

All *in vitro* and *in vivo* experiments with infectious natural isolate or rSARS-CoV-2 were conducted under appropriate biosafety level 3 (BSL3) and animal BSL3 (ABSL3) laboratories, respectively, at Texas Biomedical Research Institute. All experiments were approved by the Texas Biomed Institutional Biosafety (IBC) and Animal Care and Use Committees (IACUC).

### Cells, peptides, proteins, antibodies and viruses

African green monkey kidney epithelial cells (Vero E6, CRL-1586) were obtained from the American Type Culture Collection (ATCC, Bethesda, MD) and a Vero E6 cell line expressing hACE2 and TMPRSS2 (Vero AT) was obtained from BEI Resources (NR-54970). Cells were maintained in Dulbecco’s modified Eagle medium (DMEM) supplemented with 5% (v/v) fetal bovine serum, FBS (VWR) and 1% penicillin-streptomycin (Corning). For Vero AT, cells were treated every other passage with 5 µg/ml of puromycin for selection of AT-expressing cells.

A set of 181 peptides spanning the complete S protein of the USA-WA1/2020 strain of SARS-CoV-2 and a set of 59 peptides spanning the complete N protein of USA-WA1/2020 of SARS-CoV-2 were obtained from BEI Resources (NR-52402 and NR-52404, respectively). These peptides are 13 to 20 amino acids long, with 10 overlapping amino acids. The S2 peptide pools contain 93 peptides representing the N-terminal half of the S (MFVFLVLLPL to AEHVNNSYE) and the S2 peptide pools contain 88 peptides representing the C-terminal half of the S protein (GAEHVNNSYE to VLKGVKLHYT). Peptides were dissolved in sterile water containing 10% DMSO.

SARS-CoV-2 S protein was purchased from ACROBiosystems (Catalog# SPN-C52H9), SARS-CoV-2 E protein was purchased from ThermoFisher Scientific (Catalog# RP-87682), and SARS-CoV-2 M protein was purchased from ProteoGenix (Catalog# PX-COV-P025).

The following mouse reactive antibodies (clone, catalog number, dilution) from BioLegend, BD Biosciences, and ThermoFisher Scientific were used for analysis of T cells: CD3-PE/Cyanine7 (145-2C11, 100319, 1:400), IFN-γ-PE/Dazzle 594 (XMG1.2, 505845, 1:400), TNF-α-Brilliant Violet 785 (MP6-XT22, 506341, 1:400), IL-4-Brilliant Violet 711 (11B11, 504133, 1:100), IL-17A Alexa Flour 488 (TC11-18H10, 560221, 1:100), IL-21 Alexa Flour 647 (mhalx21, 51-7213-80, 1:100), CD4-BUV 496 (GK1.5, 612952, 1:400), CD8-BUV737 (53-6.7, 612759, 1:400), IL-10-Brilliant Violet 510 (JES5-16E3, 563277, 1:100), IL-2-PE (JES6-5H4, 12-7021-82, 1:200). The cross-reactive mouse monoclonal antibody against SARS-CoV N protein, 1C7C7, was kindly provided by Dr. Thomas Moran at the Icahn School of Medicine at Mount Sinai.

The natural isolate SARS-CoV-2 of USA-WA1/2020 (NR-52281), and α (NR-54000), β (NR-54008), δ (NR-55611), and o (NR-56461) VoC were obtained from BEI Resources. The rSARS-CoV-2 WT was generated previously ^21^, and the rSARS-CoV-2 expressing mCherryNluc (rSARS-CoV-2 mCherryNluc) was generated using the previously describe strategy ^24^.

### Reverse genetics and generation of double ORF-deficient rSARS-CoV-2

The BAC harboring the entire viral genome of SARS-CoV-2 USA-WA1/2020 strain (Accession No. MN985325) was described previously ^21^. Double deletion of accessory ORF proteins was achieved in viral fragment 1 by inverse PCR using primer pairs containing a BsaI type IIS restriction endonuclease site. All the primer sequences are available upon request. Fragments containing the double deletion of accessory ORF proteins were reassembled into the BAC using BamHI and RsrII restriction endonucleases. Virus rescues were performed as previously described ^21,36^. Viral passage 1 (P1) stocks were generated in Vero E6 cells, and then further concentrated with polyethylene glycol (System Biosciences, Catalog# LV825A-1) following the manufacturer’s protocol, and then aliquoted, titrated, and stored at −80 °C.

### Plaque assay and immunostaining

Confluent monolayers of Vero E6 cells (10^6^ cells/well, 6-well plate format, triplicates) were infected with 10-fold serial diluted viral solutions for 1 h at 37°C. After viral adsorption, cells were overlaid with post-infection media containing 1% low melting agar and incubated at 37°C. At desired times post-infection, cells were fixed overnight with 10% formaldehyde solution at 4°C. For immunostaining, cells were permeabilized with 0.5% (v/v) Triton X-100 in phosphate buffered saline (PBS) for 15 min at room temperature and immunostained using the N protein 1C7C7 monoclonal antibody (1 μg/ml) and the Vectastain ABC kit (Vector Laboratories), following the manufacturers’ instruction. After immunostaining, plates were photographed using a ChemiDoc (Bio-Rad).

### Virus passages *in vitro*

Confluent monolayers of Vero E6 cells (10^6^ cells/well, 6-well plate format, triplicates) were infected with rSARS-CoV-2 WT, Δ3a/Δ6, Δ3a/Δ7a, and Δ3a/Δ7b P1 stock at MOI 0.01. At 72 hpi, five µl of the supernatants were collected and used to infected another set of confluent Vero E6 cells (10^6^ cells/well, 6-well plate format, triplicates). After repeating the passage procedure 9 times, the resultant supernatants were labeled as P10. Then, both P1 and P10 stocks were characterized by plaque assay and immunostaining.

### RNA extraction and RT-PCR

Total RNA from mock- or virus-infected (MOI=0.01) Vero E6 cells (10^6^ cells/well, 6-well plate format) was extracted with TRIzol Reagent (Thermo Fisher Scientific) according to the manufacturer’s instructions. RT-PCR amplification of viral ORF3a, ORF6, ORF7a, ORF7b, ORF8 and N genes, was performed using Super Script II Reverse transcriptase (Thermo Fisher Scientific) and Expanded High Fidelity PCR System (Sigma Aldrich). Amplified DNA products, undigested or digested with MluI were subjected to 1.0% agarose gel analysis. All primer sequences used for RT-PCR are available upon request.

### Deep sequencing

Sequencing libraries were generated using KAPA RNA HyperPrep Kit with a 45 min adapter ligation incubation including 6 cycles of PCR with 100 ng viral RNA and 7 mM adapter. Samples were sequenced with an Illumina Hiseq X. Raw reads were quality filtered using Trimmomatic v0.39 ^37^ and mapped to a SARS-CoV-2 USA-WA1/2020 strain reference genome (Genbank Accession No. MN985325) with Bowtie2 v2.4.1 ^38^. Genome coverage was quantified with MosDepth. V0.2.6 ^39^. We genotyped each sample for low frequency variants with LoFreq* v2.1.3.1 ^40^ and filtered sites with less than 100x read depth or minor allele frequencies less than 1%. Finally, we used SnpEff v4.3t ^41^ to identify the impact of potential variants on the protein coding regions in the SARS-CoV-2 reference genome.

### Enzyme linked immunospot (ELISPOT) assay

Spleens of K18 hACE2 transgenic mice were aseptically collected at 21 dpi and minced by pressing through cell strainers. Red blood cells were removed by incubation in 0.84% ammonium chloride and, following a series of washes in RPMI 1640, cells were resuspended in RPMI 1640 supplemented with 2 mM L-glutamine, 1 mM sodium pyruvate, 10 mM HEPES, 100 U/ml penicillin, 100 μg/ml streptomycin, and 10% fetal bovine serum. Antigen-specific T cells secreting IFN-γ were enumerated using anti-mouse IFN-γ ELISPOT assay (U-Cytech, Catalogue# CT317-PB5). Cells were plated in 96 well PVDF plates at 2×10^5^ per well in duplicate and stimulated separately with SARS-CoV-2 peptide pools (2 µg/ml), Concanavalin-A (5 µg/ml; Sigma) as a positive control; or media alone as a negative control. The plates were incubated for 42-48 h and then developed according to the manufacturer’s instructions. The number of spot-forming cells (SFC) were measured using an automatic counter (Immunospot).

### Flow cytometric analysis of intracellular cytokine production

For detection of SARS-CoV-2-specific intracellular cytokine production, 10^6^ splenocytes were stimulated in 96-well round bottom plates with S1 peptide pool (5 µg/ml), or purified E (10 µg/ml) or M (1.5 µg/ml) proteins, or media alone or PMA/Ionomycin (BioLegend) as negative and positive controls, respectively, for 5 h in the presence of GolgiPlug (BD Biosciences). Following incubation, cells were surface stained for CD3, CD4, and CD8 for 30 min at 4°C, fixed and permeabilized using the cytofix/cytoperm kit (BD Biosciences), and intracellularly stained for IFN-γ, TNF-α, IL-2, IL-17A, IL-21, IL-10, and IL-4 for 30 min at room temperature. Dead cells were identified and removed using the LIVE/DEAD fixable Near-IR dead cell stain kit (Invitrogen). A positive response was defined as >3 times the background of the negative control sample. The percentage of cytokine positive cells was then calculated by subtracting the frequency of positive events in the negative control samples from that of the test samples. Events were collected on a BD LSRFortessa X-20 flow cytometer following compensation with UltraComp eBeads (Invitrogen). Data were analyzed using FlowJo v10 (Tree Star).

### Enzyme-linked immunosorbent assay (ELISA)

ELISA plates were coated with 100 ng/well of viral S protein in 50 µl of PBS overnight at 4°C. After being blocked by 2.5% BSA for 1 h at 25 °C, the plates were washed 3 times with PBS containing 0.1% Tween-20 (PBST), then incubated with 2-fold serial diluted sera at 25 °C (starting dilution of 1:50). After 2 h incubation, plates were washed 3 time with PBST, then incubated with HRP-conjugated goat anti-mouse secondary antibody (ThermoFisher Scientific, catalog# 31430) at 25°C. After 1 h, plates were washed 3 times and then developed by adding 100 µl/well of TMB-ELISA substrate (ThermoFisher Scientific, catalog# 34029). Reactions were stopped after 10 minutes by adding 50 µl/well of 3M H_2_SO_4_ to all the wells. ELISA plates were evaluated with a plate reader (Bio-Tek) at an absorbance of 450 nm.

### Plaque reduction microneutralization (PRMNT) assay

All serum samples were serially 2-fold diluted (starting dilution of 1:50) and mixed with an equal volume of DMEM containing approximately 200 PFU/well SARS-CoV-2 USA-WA1/2020 or VoC of α, β, δ, and o in a 96-well plate. After incubation at 37°C for 1 h with constant rotation, mixtures were transferred to confluent monolayers (4×10^5^ cells/well, quadruplicates) of Vero E6 cells (Vero AT cells for o), as previously described ^42^. The mixtures were replaced by fresh DMEM containing 2% FBS after incubation at 37°C for another 1 h. At 24 hpi, cells were fixed in 10% formalin solution overnight and immunostained with a SARS-CoV cross-reactive N protein monoclonal antibody 1C7C7 to visualize the plaques, as previously described ^42^. Number of plaques were measured by using an ELISPOT, and viral neutralization was quantified using a sigmoidal dose response curve. Mock-infected cells and SARS-CoV-2-infected cells in the absence of serum were included as internal controls. Neutralizing titer at 50% inhibition (NT_50_) was calculated for each serum sample.

### Evaluation of lung pathological lesions

Macroscopic pathology scoring was evaluated using ImageJ software to determine the percentage of the total surface area of the lung (dorsal and ventral view) affected by consolidation, congestion, and atelectasis, as previously described ^43^.

### Multiplex cytokines and chemokines assay

Multiple cytokines and chemokines (IFN-α, IFN-γ, IL-6, IL-10, IL-17A, MCP-1, RANTES, and TNF-α) were measured using a custom 8-plex panel mouse ProcartaPlex assay (ThermoFisher Scientific, catalog# PPX-08-MXGZGFX), following the manufacturer’s instructions. The assay was performed in a BSL3 laboratory and samples were decontaminated by an overnight incubation in 10% formaldehyde solution before readout on a Luminex 100/200 System running on Xponent v4.2 with the following parameters: gate 5,000-25,000, 50 μl of sample volume, 50 events per bead, sample timeout 120 s, low PMT (LM×100/200: Default). Acquired data were analyzed using ProcartaPlex Analysis Software v1.0.

### Animal experiments

All animal protocols were approved by Texas Biomedical Research Institute IACUC. Five-week-old female K18 hACE2 transgenic mice were purchased from The Jackson Laboratory and five-week-old female golden Syrian hamsters were purchased from Charles River Laboratories. All the animals were maintained in the animal facility at Texas Biomedical Research Institute under specific pathogen-free conditions. Animal infection was performed by intranasally inoculation after being anesthetized following gaseous sedation in an isoflurane chamber.

#### Pathogenicity analysis of double ORF-deficient rSARS-CoV-2 in K18 hACE2 transgenic mice

Five-week-old K18 hACE2 transgenic mice (n=8) were intranasally (IN) infected (2×10^5^ PFU) with the double ORF-deficient rSARS-CoV-2 or rSARS-CoV-2 WT. At 2 and 4 dpi, four mice per group were humanely sacrificed to collect lungs and nasal turbinates, and gross images of lungs were taken using an iPhone 6s (Apple). Then, nasal turbinates and lungs were homogenized and processed as previously described ^22^. Tissue homogenates were centrifuged at 12,000x g for 5 min at 4°C, and clarified supernatant were collected for further measurement of viral titers and cytokine and chemokine induction. Another set of five-week-old K18 hACE2 transgenic mice (n=5) were mock infected or infected (IN, 2×10^5^ PFU) with double ORF-deficient rSARS-CoV-2 or rSARS-CoV-2 WT to evaluate body weight changes and survival rate daily for 21 days. The surviving mice were bled to collect serum to assess total IgG levels against viral full-length S protein. Afterward, mice were sacrificed to collect spleens for analysis of T cell responses in splenocytes.

#### Immunization with rSARS-CoV-2 Δ3a/Δ7b protected K18 hACE2 transgenic mice from lethal challenge of rSARS-CoV-2 mCherryNluc

Five-week-old K18 hACE2 transgenic mice (n=8) were mock-vaccinated or vaccinated with rSARS-CoV-2 Δ3a/Δ7b (IN; 2×10^5^ PFU). At 21 days post-vaccination, all vaccinated mice were IN challenged with rSARS-CoV-2 mCherryNluc (10^5^ PFU). At 2 and 4 days post-challenge, mice (n=4) were anesthetized with isoflurane and imaged immediately under an *in vivo* imaging system (IVIS, AMI HTX) after retro-orbitally injected with 100 µl of Nano-Glo luciferase substrate (Promega). The bioluminescence data acquisition and analysis were performed using the Aura program (Spectral Imaging Systems). Flux measurements were acquired from the region of interest. Then, lungs were excised to analyze mCherry expression under the IVIS, and brightfield images were taken using an iPhone 6s for pathological lesion analysis. Finally, nasal turbinates and lungs were homogenized as described above and the clarified supernatants of homogenate were collected for further measurement of viral titers, Nluc activity, and cytokine and chemokine induction. Another set of five-week-old K18 hACE2 transgenic mice (n=5) were mock-vaccinated or vaccinated (IN; 2×10^5^ PFU) with rSARS-CoV-2 Δ3a/Δ7b and challenged (IN; 10^5^ PFU) with rSARS-CoV-2 mCherryNluc at 21 days post-vaccination. Body weight and survival rate were assessed by monitoring daily for 15 days post-challenge.

#### Immunization of hamsters with double ORF-deficient rSARS-CoV-2 prevented viral replication and shedding

Five-week-old golden Syrian hamsters (n=4) were mock vaccinated or vaccinated (IN; 4×10^5^ PFU) with the double ORF-deficient rSARS-CoV-2. Mock-vaccinated and vaccinated hamsters were challenged (IN; 2×10^5^ PFU) with rSARS-CoV-2 mCherryNluc at 21 days post-vaccination and then housed with non-treated susceptible contact hamsters (n=4). At 2 and 4 days post-challenge, all hamsters were anesthetized and imaged immediately after intraperitoneal injection of 200 µl of Nano-Glo luciferase substrate. At 4 days post-challenge, all hamsters were euthanized and their lungs were excised. Reporter mCherry expression in the lungs were evaluated by IVIS as previously described ^44^. Thereafter, all lungs and nasal turbinates were collected and homogenized in 2 ml of PBS, and viral titers and Nluc activity were determined in the clarified supernatant of lung and nasal turbinate homogenate ^44^.

#### Immunization of hamsters with double ORF-deficient rSARS-CoV-2 prevented viral transmission

Five-week-old golden Syrian hamsters (n=4) were mock-vaccinated or vaccinated (IN; 4×10^5^ PFU) with the double ORF-deficient rSARS-CoV-2. At 18 days post-vaccination, all hamsters were bled and sera were isolated for evaluation of neutralization capacity by PRMNT assay, as previously described ^42^. At 21 days post-vaccination, mock-vaccinated and vaccinated hamsters were housed with donor hamsters, which had been infected (IN; 2×10^5^ PFU) with rSARS-CoV-2 mCherryNluc for 24 h. At 2 and 4 dpi with rSARS-CoV-2 mCherryNluc, all hamsters were anesthetized and imaged immediately after intraperitoneal injection of 200 µl of Nano-Glo luciferase substrate. At 4 dpi with rSARS-CoV-2 mCherryNluc, all hamsters were euthanized and their lungs were excised. Reporter mCherry expression in the lungs of hamsters were evaluated under an IVIS, as previously described above. Thereafter, all lungs and nasal turbinates were collected and homogenized in 2 ml of PBS, and viral titers and Nluc activity were determined in the clarified supernatant of lung and nasal turbinate homogenate, as previously described above.

### Statistical analysis

Data representative of three independent experiments in triplicate have been used. All data represent the mean ± standard deviation (SD) for each group and analyzed by SPSS13.0 (IBM). A two-tailed Student’s t-test was used to compare the mean between two groups. *P* values less than 0.05 (*P*<0.05) were considered statistically significant.

## Acknowledgements

Funding for this work provided by the National Institutes of Health 1R01AI161175 to J.J.K., L.M.-S., and M.R.W., R01 AI145332 to J.J.K., and L.M.-S., and the Center for Research on Influenza Pathogenesis and Transmission (CRIPT), a NIAID-funded Center of Excellence for Influenza Research and Response (CEIRR, contract # 75N93021C00014) to L.M.-S.

## Author contributions

C.Y. and L.M.-S. conceived the study. C.Y., J.-G.P., and K.C. performed the majority of experiments. P.D. and A.K. analyzed the T cell response. A.A.-G., A.G.-V., and J.B.T. evaluated the expression of cytokines and chemokines. M.R.W., J.J.K., and R.K.P. provided critical reagents. C.Y. and L.M.-S. drafted the manuscript.

## Data Availability

All the recombinant viruses described in the study are available at the following website: https://www.txbiomed.org/services-2/virus-request/.

## Extended Data

**Extended Data Figure 1. Generation of double ORF-deficient rSARS-CoV-2**.

**(A)** Schematic representations of the double ORF-deficient rSARS-CoV-2 genomes (no drawn to scale).

**(B)** Confirmation of the double ORF-deficient rSARS-CoV-2 by RT-PCR amplification of ORF3a, ORF6, ORF7a, ORF7b and N.

**(C)** Deep sequencing analysis of the double ORF-deficient rSARS-CoV-2 genome. Non-reference alleles present in less than 10% of reads are not shown. Amino acid changes respective to rSARS-CoV-2 WT are indicated.

**Extended Data Figure 2. Induction of cytokines and chemokines in lung homogenates, and activation of cytokine positive CD4 and CD8 splenocytes against viral components upon double ORF-deficient rSARS-CoV-2 infection**.

**(A)** Cytokine and chemokine levels were measured in the lung homogenates of K18 hACE2 transgenic mice infected (2×10^5^ PFU/mouse) with WT or double ORF-deficient rSARS-CoV-2 at 2 and 4 dpi. *, *P*<0.05; **, *P*<0.01.

**(B)** Intracellular cytokine positive CD4^+^ T cells in the splenocytes of the double ORF-deficient rSARS-CoV-2-infected K18 hACE2 transgenic mice were analyzed after stimulation of S1 peptide pool, E, and M using flow cytometry. The splenocytes from the two surviving K18 hACE2 transgenic mice infected with rSARS-CoV-2 WT (n=5) for 21 days were collected as a positive control.

**(C)** Intracellular cytokine positive CD8^+^ T cells in the splenocytes of the double ORF-deficient rSARS-CoV-2-infected K18 hACE2 transgenic mice were analyzed after stimulation of S1 peptide pool, E, and M using flow cytometry. The splenocytes from the two surviving K18 hACE2 transgenic mice infected with rSARS-CoV-2 WT (n=5) for 21 days were collected as a positive control.

**Extended Data Figure 3. Analysis of mCherry expression, pathological lesions, viral replications, and cytokine and chemokines induction in the lungs of mock-or rSARS-CoV-2 Δ3a/Δ7b-vaccinated K18 hACE2 transgenic mice at 2 and 4 days post-challenge with rSARS-CoV-2 mCherryNluc**.

**(A)** Expression of mCherry in the lungs of K18 hACE2 transgenic mice challenged with rSARS-CoV-2 mCherryNluc.

**(B)** Quantitative analysis of mCherry intensity in the lungs of challenged K18 hACE2 transgenic mice.

**(C)** Pathological lesions on the lungs surface of K18 hACE2 transgenic mice challenged with rSARS-CoV-2 mCherryNluc.

**(D)** Quantitative analysis of pathological lesion on the lungs surface of K18 hACE2 transgenic mice challenged with rSARS-CoV-2 mCherryNluc.

**(E)** Viral replication in the lungs and nasal turbinates of K18 hACE2 transgenic mice at 2 and 4 days post-challenge with rSARS-CoV-2 mCherryNluc.

**(F)** Nluc activity in the clarified lung and nasal turbinate homogenates of the K18 hACE2 transgenic mice was determined at 2 and 4 days post-challenge with rSARS-CoV-2 mCherryNluc.

**(G)** Cytokine and chemokine levels were measured in the lung homogenates of K18 hACE2 transgenic mice at 2 and 4 days post-challenge with rSARS-CoV-2 mCherryNluc. *, *P*<0.05; **, *P*<0.01; ns, not significant.

**Extended Data Figure 4. Analysis of the mCherry expression, viral replication, and Nluc activity in the lungs of the double ORF-deficient rSARS-CoV-2-vaccinated hamsters challenged and co-housed with susceptible contact hamsters**.

**(A)** Expression of mCherry in the lungs of challenged and contact hamsters.

**(B)** Quantitative analysis of mCherry intensity in the lungs of challenged and contact hamsters.

**(C)** Replication of rSARS-CoV-2 mCherryNluc in the lungs and nasal turbinates of challenged and contact hamsters.

**(D)** Nluc activity in the clarified lung and nasal turbinate homogenate of infected and contact hamsters.

**Extended Data Figure 5. Analysis of the mCherry expression, virus replication, and Nluc activity in the lungs of rSARS-CoV-2 mCherryNluc-infected donor hamsters and their contacts**.

**(A)** Expression of mCherry in the lungs of infected donor and vaccinated contact hamsters.

**(B)** Quantitative analysis of mCherry intensity in the lungs of infected donor and vaccinated contact hamsters.

**(C)** Replication of rSARS-CoV-2 mCherryNluc in the lungs and nasal turbinates of infected donor and vaccinated contact hamsters.

**(D)** Nluc activity in the clarified lung and nasal turbinate homogenates of infected donor and vaccinated contact hamsters.

## Notes

### Competing Interest Statement

The authors have declared no competing interest.

## REFERENCES

1 Moriyama, M., Hugentobler, W. J. & Iwasaki, A. Seasonality of Respiratory Viral Infections. Annu Rev Virol 7, 83–101, doi:10.1146/annurev-virology-012420-022445 (2020).

2 Cui, J., Li, F. & Shi, Z. L. Origin and evolution of pathogenic coronaviruses. Nat Rev Microbiol 17, 181–192, doi:10.1038/s41579-018-0118-9 (2019).

3 Yao, H. P. et al. Molecular Architecture of the SARS-CoV-2 Virus. Cell 183, 730–738, doi:10.1016/j.cell.2020.09.018 (2020).

4 Hu, B., Guo, H., Zhou, P. & Shi, Z. L. Characteristics of SARS-CoV-2 and COVID-19. Nat Rev Microbiol 19, 141–154, doi:10.1038/s41579-020-00459-7 (2021).

5 Ren, Y. J. et al. The ORF3a protein of SARS-CoV-2 induces apoptosis in cells. Cell Mol Immunol 17, 881–883, doi:10.1038/s41423-020-0485-9 (2020).

6 Miorin, L. et al. SARS-CoV-2 Orf6 hijacks Nup98 to block STAT nuclear import and antagonize interferon signaling. P Natl Acad Sci USA 117, 28344–28354, doi:10.1073/pnas.2016650117 (2020).

7 Zhou, Z. L. et al. Structural insight reveals SARS-CoV-2 ORF7a as an immunomodulating factor for human CD14(+) monocytes. Iscience 24, doi:10.1016/j.isci.2021.102187 (2021).

8 Shemesh, M. et al. SARS-CoV-2 suppresses IFNbeta production mediated by NSP1, 5, 6, 15, ORF6 and ORF7b but does not suppress the effects of added interferon. PLoS Pathog 17, e1009800, doi:10.1371/journal.ppat.1009800 (2021).

9 Park, M. D. Immune evasion via SARS-CoV-2 ORF8 protein? Nat Rev Immunol 20, 408, doi:10.1038/s41577-020-0360-z (2020).

10 Corbett, K. S. et al. SARS-CoV-2 mRNA vaccine design enabled by prototype pathogen preparedness. Nature 586, 567–571, doi:10.1038/s41586-020-2622-0 (2020).

11 Polack, F. P. et al. Safety and Efficacy of the BNT162b2 mRNA Covid-19 Vaccine. New Engl J Med 383, 2603–2615, doi:10.1056/NEJMoa2034577 (2020).

12 Kustin, T. et al. Evidence for increased breakthrough rates of SARS-CoV-2 variants of concern in BNT162b2-mRNA-vaccinated individuals. Nat Med 27, 1379-+, doi:10.1038/s41591-021-01413-7 (2021).

13 Choi, A. et al. Serum Neutralizing Activity of mRNA-1273 against SARS-CoV-2 Variants. J Virol 95, doi:10.1128/JVI.01313-21 (2021).

14 Barrett, A. D. T. Yellow fever live attenuated vaccine: A very successful live attenuated vaccine but still we have problems controlling the disease. Vaccine 35, 5951–5955, doi:10.1016/j.vaccine.2017.03.032 (2017).

15 Stokes, A., Bauer, J. H., Hudson, N. P. & Mortimer, P. P. The transmission of yellow fever to Macacus rhesus (Reprint from JAMA, vol 96, pg 253-254, 1928). Rev Med Virol 11, 141–148, doi:DOI 10.1002/rmv.311 (2001).

16 Pulendran, B. Learning immunology from the yellow fever vaccine: innate immunity to systems vaccinology. Nat Rev Immunol 9, 741–747, doi:10.1038/nri2629 (2009).

17 Johnson, B. A. et al. Loss of furin cleavage site attenuates SARS-CoV-2 pathogenesis. Nature 591, 293–299, doi:10.1038/s41586-021-03237-4 (2021).

18 Wang, Y. et al. Scalable live-attenuated SARS-CoV-2 vaccine candidate demonstrates preclinical safety and efficacy. Proc Natl Acad Sci U S A 118, doi:10.1073/pnas.2102775118 (2021).

19 Trimpert, J. et al. Development of safe and highly protective live-attenuated SARS-CoV-2 vaccine candidates by genome recoding. Cell Rep 36, doi:10.1016/j.celrep.2021.109493 (2021).

20 Silvas, J. A. et al. Contribution of SARS-CoV-2 Accessory Proteins to Viral Pathogenicity in K18 Human ACE2 Transgenic Mice. J Virol 95, doi:10.1128/JVI.00402-21 (2021).

21 Ye, C. et al. Rescue of SARS-CoV-2 from a Single Bacterial Artificial Chromosome. mBio 11, e0216820, doi:10.1128/mBio.02168-20 (2020).

22 Oladunni, F. S. et al. Lethality of SARS-CoV-2 infection in K18 human angiotensin-converting enzyme 2 transgenic mice. Nat Commun 11, 6122, doi:10.1038/s41467-020-19891-7 (2020).

23 Hassan, A. O. et al. A SARS-CoV-2 Infection Model in Mice Demonstrates Protection by Neutralizing Antibodies. Cell 182, 744–753 e744, doi:10.1016/j.cell.2020.06.011 (2020).

24 Ye, C. J. et al. Analysis of SARS-CoV-2 infection dynamic in vivo using reporter-expressing viruses. P Natl Acad Sci USA 118, doi:10.1073/pnas.2111593118 (2021).

25 Monto, A. S. The Future of SARS-CoV-2 Vaccination - Lessons from Influenza. N Engl J Med 385, 1825–1827, doi:10.1056/NEJMp2113403 (2021).

26 McCray, P. B., Jr. et al. Lethal infection of K18-hACE2 mice infected with severe acute respiratory syndrome coronavirus. J Virol 81, 813–821, doi:10.1128/JVI.02012-06 (2007).

27 Winkler, E. S. et al. SARS-CoV-2 infection of human ACE2-transgenic mice causes severe lung inflammation and impaired function. Nat Immunol 21, 1327–1335, doi:10.1038/s41590-020-0794-2 (2020).

28 Israelow, B. et al. Adaptive immune determinants of viral clearance and protection in mouse models of SARS-CoV-2. Sci Immunol 6, doi:10.1126/sciimmunol.abl4509 (2021).

29 Lu, S. Heterologous prime-boost vaccination. Curr Opin Immunol 21, 346–351, doi:10.1016/j.coi.2009.05.016 (2009).

30 Kardani, K., Bolhassani, A. & Shahbazi, S. Prime-boost vaccine strategy against viral infections: Mechanisms and benefits. Vaccine 34, 413–423, doi:10.1016/j.vaccine.2015.11.062 (2016).

31 Winkler, E. S. et al. SARS-CoV-2 causes lung infection without severe disease in human ACE2 knock-in mice. J Virol 96, e01511–01521 (2021).

32 Rapeport, G. et al. SARS-CoV-2 Human Challenge Studies - Establishing the Model during an Evolving Pandemic. New Engl J Med 385, 961–964, doi:10.1056/NEJMp2106970 (2021).

33 Sia, S. F. et al. Pathogenesis and transmission of SARS-CoV-2 in golden hamsters. Nature 583, 834–838, doi:10.1038/s41586-020-2342-5 (2020).

34 Imai, M. et al. Syrian hamsters as a small animal model for SARS-CoV-2 infection and countermeasure development. P Natl Acad Sci USA 117, 16587–16595, doi:10.1073/pnas.2009799117 (2020).

35 Zhu, N. et al. A Novel Coronavirus from Patients with Pneumonia in China, 2019. N Engl J Med 382, 727–733, doi:10.1056/NEJMoa2001017 (2020).

36 Chiem, K., Ye, C. & Martinez-Sobrido, L. Generation of Recombinant SARS-CoV-2 Using a Bacterial Artificial Chromosome. Curr Protoc Microbiol 59, e126 (2020).

37 Bolger, A. M., Lohse, M. & Usadel, B. Trimmomatic: a flexible trimmer for Illumina sequence data. Bioinformatics 30, 2114–2120, doi:10.1093/bioinformatics/btu170 (2014).

38 Langmead, B. & Salzberg, S. L. Fast gapped-read alignment with Bowtie 2. Nat Methods 9, 357–359, doi:10.1038/Nmeth.1923 (2012).

39 Pedersen, B. S. & Quinlan, A. R. Mosdepth: quick coverage calculation for genomes and exomes. Bioinformatics 34, 867–868, doi:10.1093/bioinformatics/btx699 (2018).

40 Wilm, A. et al. LoFreq: a sequence-quality aware, ultra-sensitive variant caller for uncovering cell-population heterogeneity from high-throughput sequencing datasets. Nucleic Acids Res 40, 11189–11201, doi:10.1093/nar/gks918 (2012).

41 Cingolani, P. et al. A program for annotating and predicting the effects of single nucleotide polymorphisms, SnpEff: SNPs in the genome of Drosophila melanogaster strain w(1118); iso-2; iso-3. Fly 6, 80–92, doi:10.4161/fly.19695 (2012).

42 Park, J. G. et al. Rapid in vitro assays for screening neutralizing antibodies and antivirals against SARS-CoV-2. J Virol Methods 287, 113995, doi:10.1016/j.jviromet.2020.113995 (2021).

43 Jensen, E. C. Quantitative Analysis of Histological Staining and Fluorescence Using ImageJ. Anat Rec 296, 378–381, doi:10.1002/ar.22641 (2013).

44 Chiem, K. et al. A Bifluorescent-Based Assay for the Identification of Neutralizing Antibodies against SARS-CoV-2 Variants of Concern In Vitro and In Vivo. J Virol 95, e0112621, doi:10.1128/JVI.01126-21 (2021).

