## Supplementary figures and images for "Immunization with recombinant accessory protein-deficient SARS-CoV-2 protects against lethal challenge and viral transmission"

### Supplementary Figures

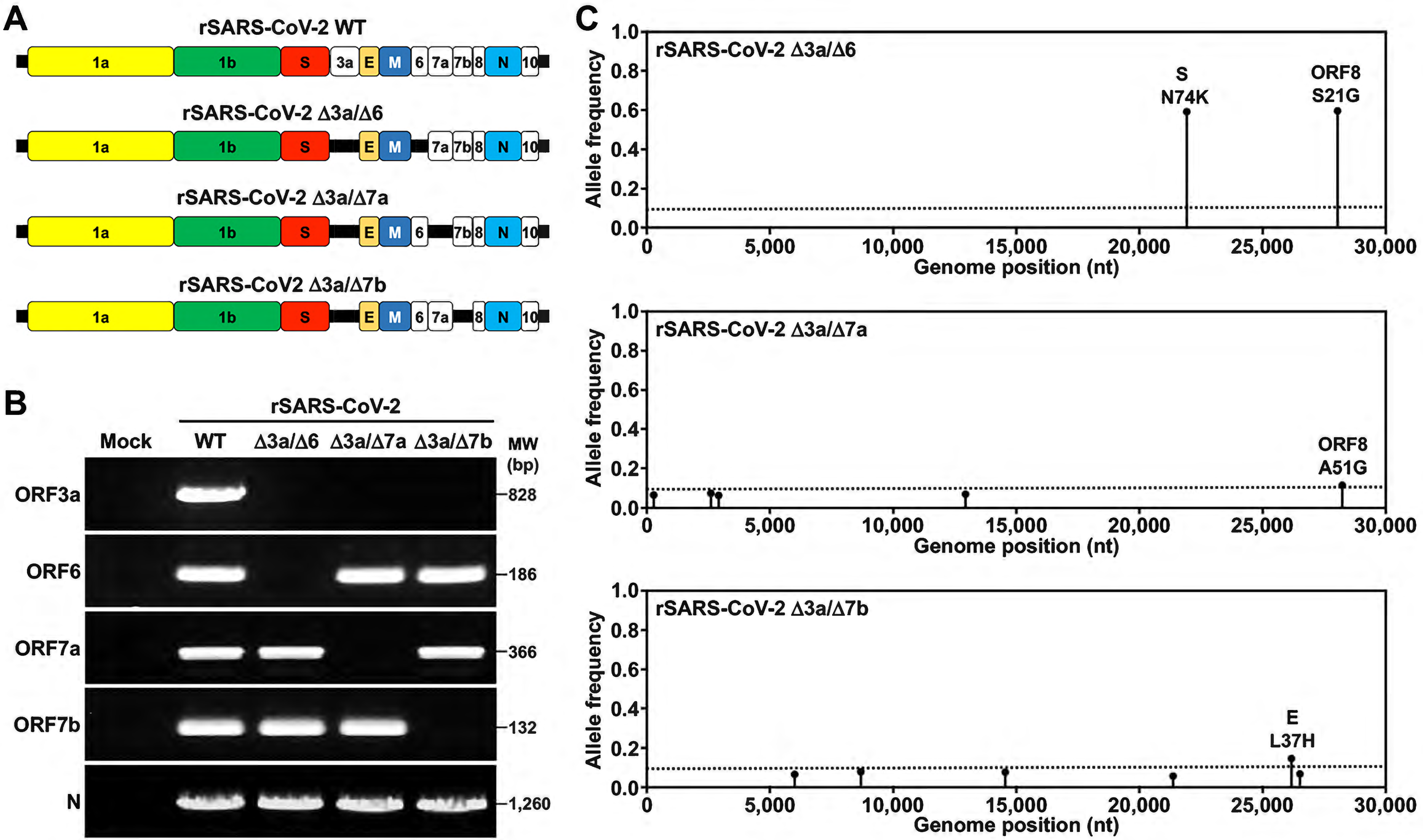

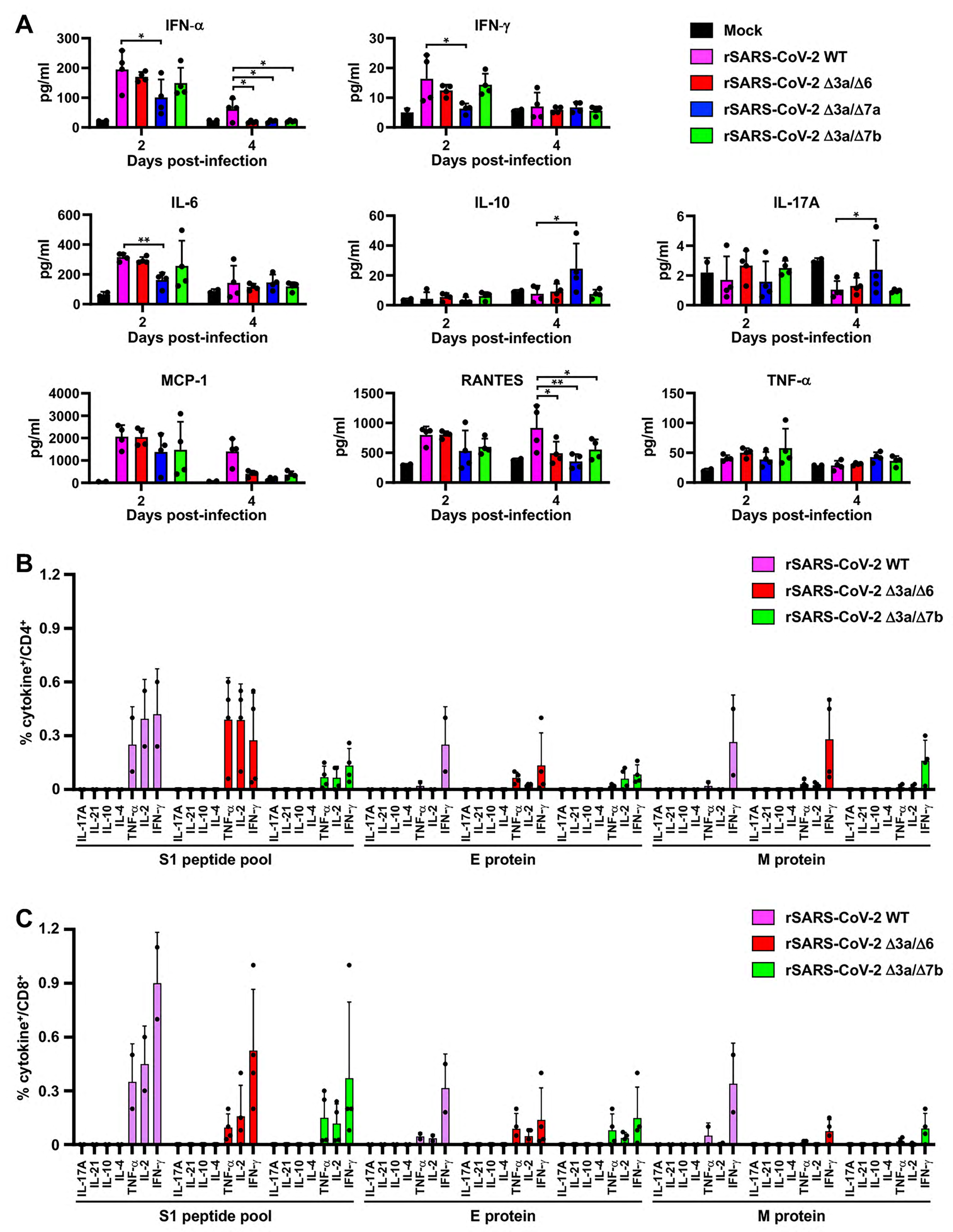

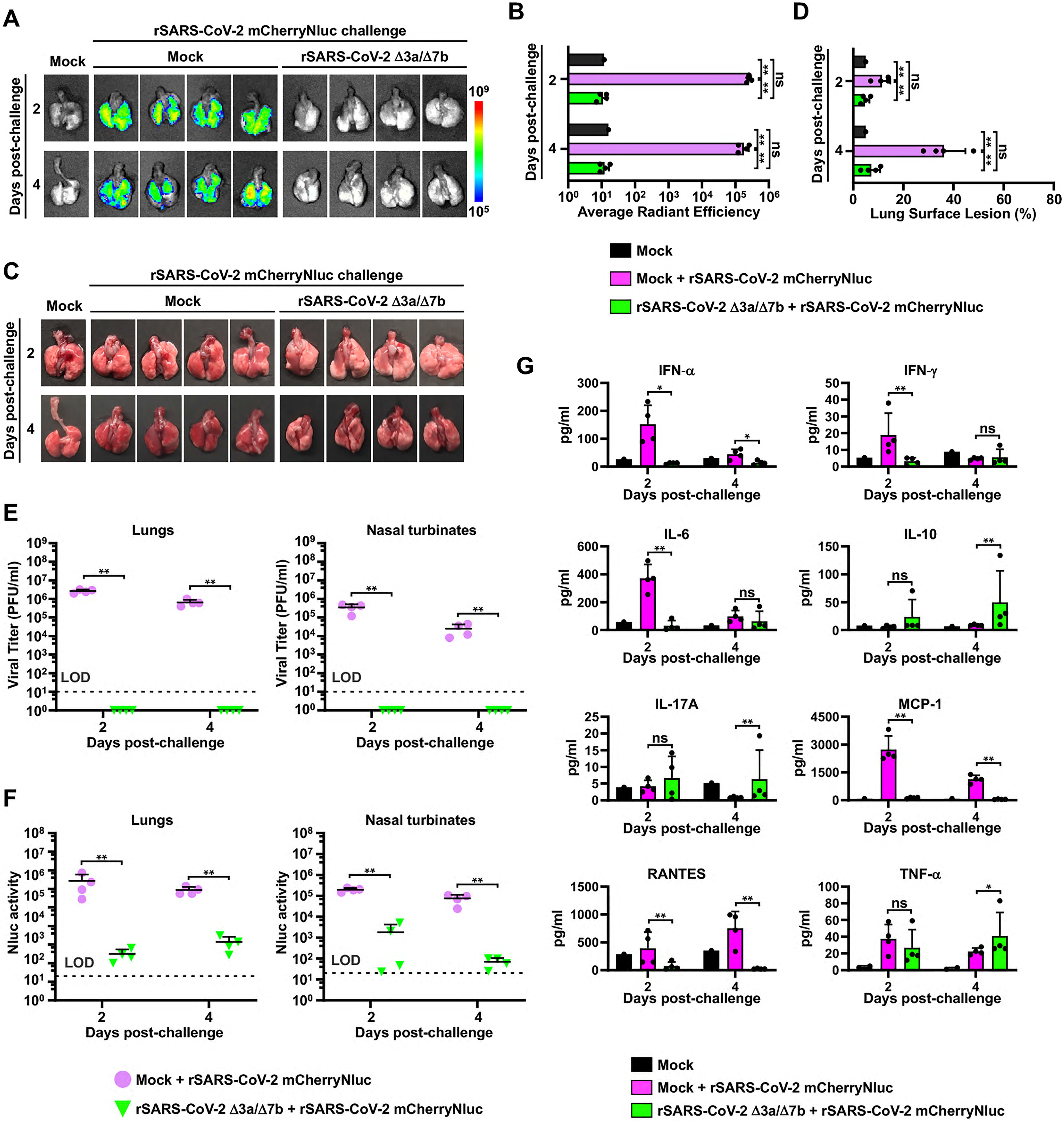

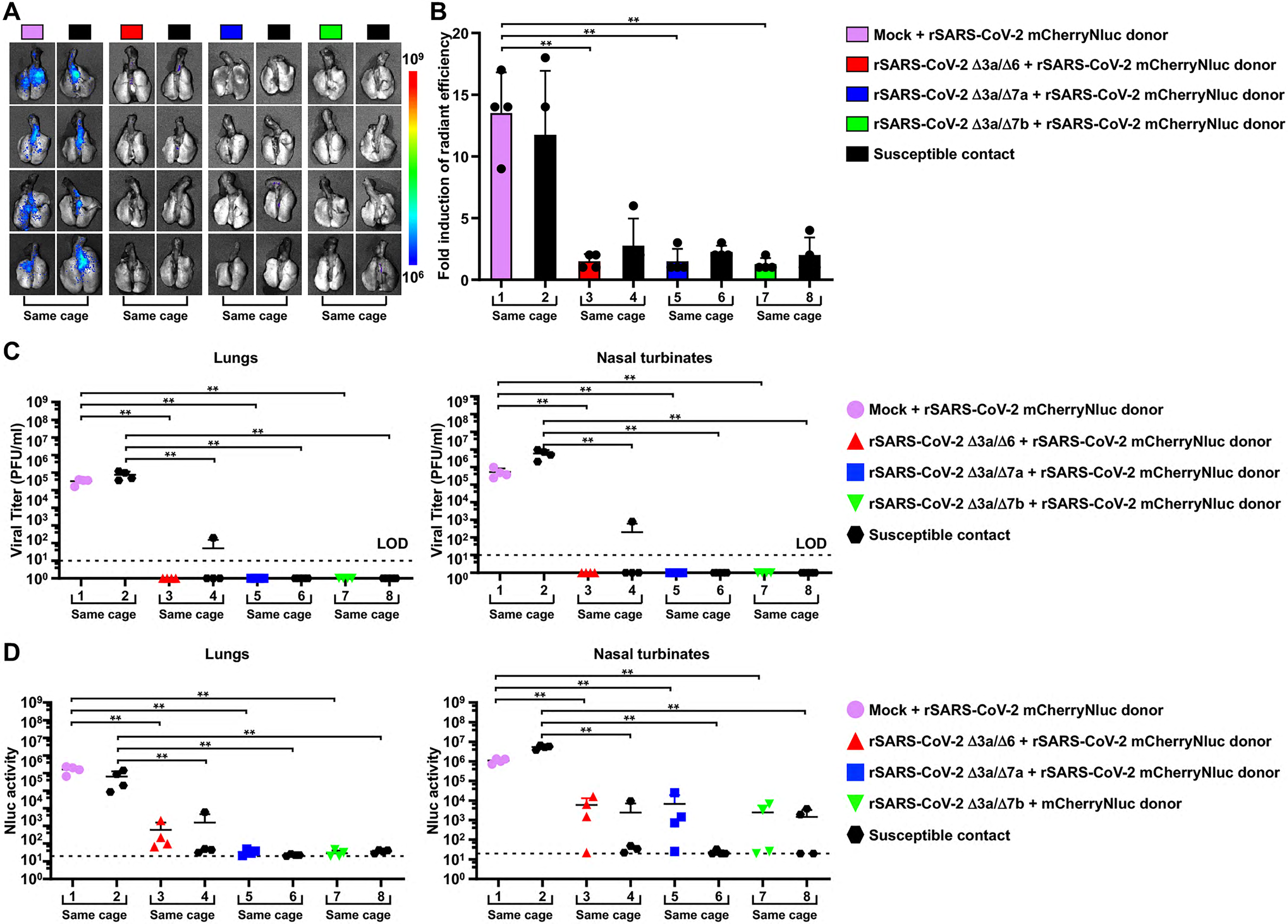

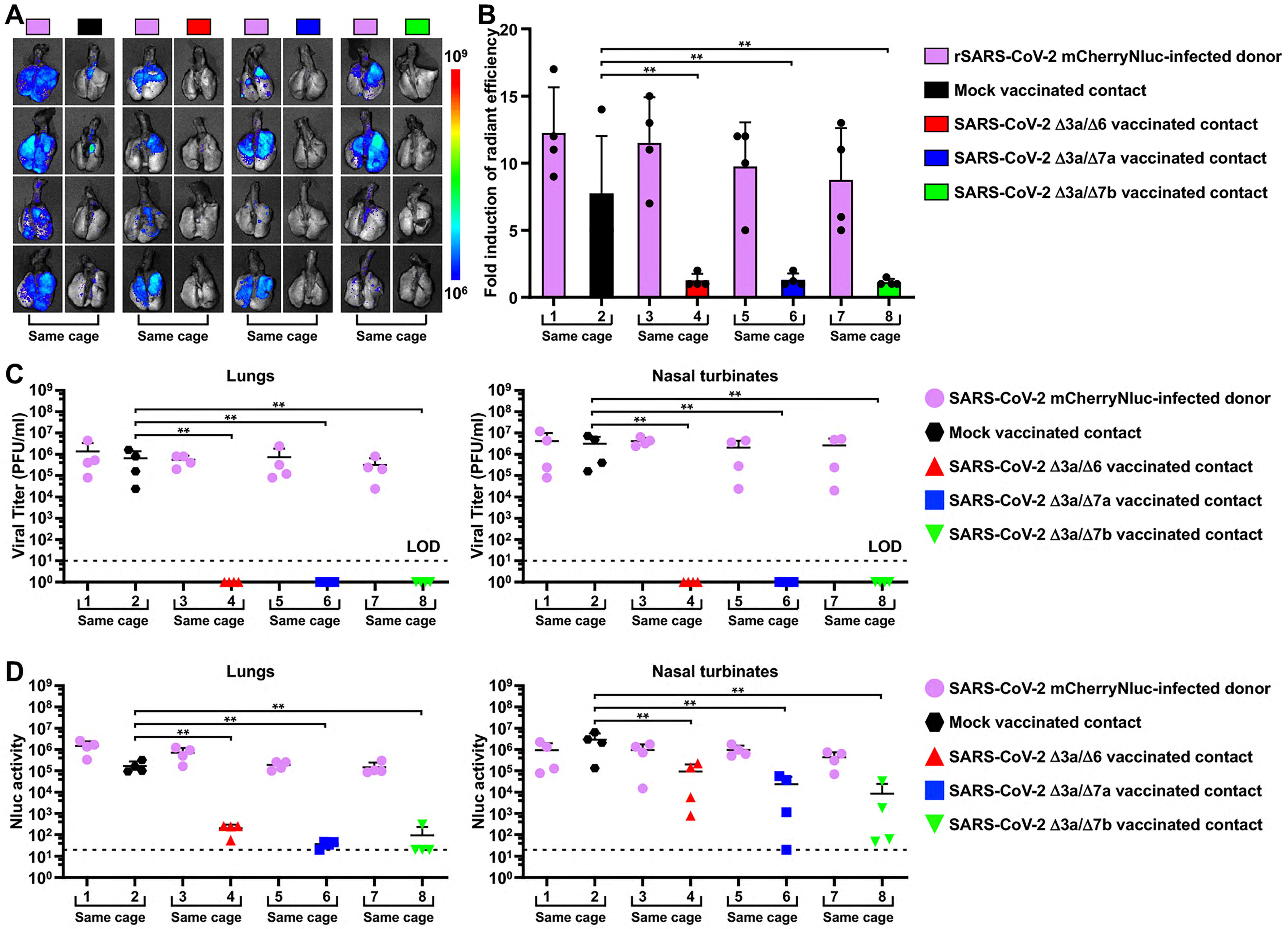
